# S-nitrosylation-mediated unfolded protein response maintains hematopoietic progenitors

**DOI:** 10.1101/2023.06.30.547214

**Authors:** Bumsik Cho, Mingyu Shin, Eunji Chang, Seogho Son, Jiwon Shim

## Abstract

The *Drosophila* lymph gland houses blood progenitors that give rise to myeloid-like blood cells. Initially, blood progenitors proliferate, but later become quiescent to maintain multipotency before differentiation. Despite the identification of various factors involved in multipotency maintenance, the cellular mechanism regulating blood progenitor quiescence remains elusive. Here, we reveal the expression of nitric oxide synthase in blood progenitors, generating nitric oxide for post-translational S-nitrosylation of protein cysteine residues. S-nitrosylation activates the Ire1-Xbp1-mediated unfolded protein response, leading to G2 cell cycle arrest. Specifically, we identify the epidermal growth factor receptor as a target of S-nitrosylation, resulting in its retention within the endoplasmic reticulum and blockade of its receptor function. Collectively, our findings highlight developmentally programmed S-nitrosylation as a critical mechanism that induces protein quality control in blood progenitors, maintaining their undifferentiated state by inhibiting cell cycle progression and rendering them unresponsive to paracrine factors.

## Introduction

Nitric oxide (NO) is a gaseous signaling molecule involved in a variety of biological processes, including neurotransmission, cardiovascular function, and immune responses^1–4^. NO is produced by nitric oxide synthase (Nos), which utilizes L-arginine as a substrate and oxygen and NADPH as co-substrates to synthesize NO^5, 6^. Nos contains two functional domains: reductase and oxygenase domains; the reductase domain donates electrons to the oxygenase domain, which uses them to generate NO^5,7,8^. NO can either function locally, near its site of synthesis, or it can diffuse across membranes to trigger paracrine signaling pathways in distant cells^9^. There are multiple modes of NO activation. NO can activate classical signaling pathways mediated by soluble guanylyl cyclase (sGC) family proteins, generating cGMP as a second messenger to induce the cGMP-dependent protein kinase (PKG) in cellular responses^10–12^. In addition, NO reacts with specific amino acid residues in proteins or binds transition metals to transfer NO molecules to neighboring metal complexes^13, 14^. Finally, NO can also post-translationally modify the thiol groups of cysteine residues in a process named S-nitrosylation, a major cGMP-independent NO signaling mechanism that modifies protein structure and function^15, 16^. S-nitrosylation targets thousands of proteins in multiple biological processes, and its dysregulation is implicated in various diseases^17, 18^. For example, caspase-3 is a representative target of S-nitrosylation, which inhibits its cleavage under normal conditions but promotes it during programmed cell death^17,19–21^.

The unfolded protein response (UPR) is a cellular protein quality control pathway that protects cells from accumulation of unfolded or misfolded proteins, particularly during stress conditions, such as aging, starvation, or neurodegenerative disorders^22–25^. Its primary function is the mitigation of a buildup of misfolded proteins by halting translation, degrading unfolded proteins, and restoring appropriate protein folding with the help of molecular chaperones. Three representative pathways are activated during UPR: the Ire1–Xbp1, PERK/Atf4, and Atf6 pathways^26^. Inositol-requiring enzyme 1 (Ire1) is an ER-tethered ribonuclease and sensor for unfolded proteins in the ER. It is activated through oligomerization and autophosphorylation when ER stress is sensed, and this leads to alternative splicing of inactive *Xbp1* mRNA into an active form. The spliced *Xbp1* encodes a bZIP transcription factor that activates transcription of chaperones, degradation of ER-associated proteins, and/or lipid biogenesis, each of which is critical for suppressing the UPR^27, 28^. PERK is a type I transmembrane serine/threonine kinase, also phosphorylated by ER stress; it attenuates protein translation by inhibiting eIF2α or through the selective translation of *Atf4* transcripts. In turn, Atf4 activates the transcription of target genes involved in apoptosis or protein translation^25, 29^. Atf6 is an ER-resident type II transmembrane protein with a bZIP domain at its N-terminus^30, 31^. During ER stress, Atf6 is translocated from the ER to the Golgi, where it is cleaved to release a bZIP domain transcription factor that induces transcription of *Xbp1* and chaperones that are critical regulators of the UPR^31, 32^.

The *Drosophila* lymph gland is a larval hematopoietic organ that houses blood progenitor cells, including their descendant mature blood cells and the microenvironment niche. Blood cells in the lymph gland are classified based on their markers: the medullary zone (MZ), the cortical zone (CZ), and the posterior signaling center (PSC)^33–35^. The MZ contains a group of blood progenitor cells in its inner core; they are medially localized adjacent to the dorsal vessel. Progenitor cells proliferate during the early instar stage but later cease their division and give rise to mature blood cells from the distal margins^36–39^. Mature blood cells that arise from blood progenitors placed close to the outer boundary of the lymph gland, farthest away from the dorsal vessel, belong to the CZ. Between the progenitor and mature blood cells is the intermediate zone (IZ), which contains independent cell types with some, but not all, of the markers common in the MZ and CZ^35, 40^. When unchallenged by infection or stress, the lymph gland undergoes a stereotypical developmental program regulated either by local signals propagated from its microenvironment—the posterior signaling center and the dorsal vessel, by cell autonomous factors downstream of the Wg, Hh, and JAK/STAT pathways^41–45^, or by systemic signals originating from other tissues or the environment^46–48^. Multiple markers to label the various cell types in the lymph gland have been validated^49–51^. The most prominent of these is *Dome^Meso^*, a marker representative of the entire progenitor population, including differentiating progenitors^52^, while *Tep4*^53, 54^, and low-level collier protein expression (col^Low^), mark early core progenitors^50, 55^. Differentiating blood cells co-express the progenitor marker *Dome^Meso^* and the earliest differentiating blood cell marker, *Hml* or Pxn^35,40,50^. Cells of the CZ express markers for differentiated blood cells: plasmatocytes, crystal cells, and lamellocytes. Plasmatocytes are evolutionarily related to mammalian macrophages^56^ and account for about 95% of differentiated blood cells; initially, they express *Hml* or Pxn, which subsequently activate NimC1 upon maturation^57^. Crystal cells, which are analogous to platelets, function in wound response and melanization. Under steady-state conditions, approximately 50–100 crystal cells develop in a typical lymph gland lobe and are specifically marked by hindsight (hnt) or lozenge (lz)^58, 59^. Lamellocytes are rarely observed under normal growth conditions because they differentiate in response to active immune challenges or stress^60, 61^. Complementing above markers, recent single-cell RNA sequencing techniques have identified novel markers and more heterogeneity within lymph gland cells among populations than was evident from classical genetic analyses^62–65^.

The epidermal growth factor receptor (EGFR) is an evolutionarily conserved receptor tyrosine kinase that plays a crucial role in many aspects of tissue development and homeostasis. Although primarily associated with epithelial tissue growth during animal development, EGFR has been increasingly recognized for its inappropriate activation in tumorigenesis and as a target for anticancer drugs currently used in clinical practices. In *Drosophila*, there is only one EGFR that controls cell proliferation, migration, survival, and cell fate determination in tissues, including the gut, discs, reproductive organs, and the lymph gland^66–71^. The *Drosophila* EGFR is activated by one of three ligands: spitz (spi), Keren (Krn), or gurken (grk). These ligands are expressed as membrane-tethered precursors and are cleaved by an intramembrane protease, rhomboid (rho)^72^, in a tissue-or context-dependent manner. Upon ligand binding, the intrinsic tyrosine kinase domain of EGFR is phosphorylated, which triggers signaling cascades, and rapidly internalized by endosomes to downregulate its receptor function^73^. Consequently, the combination of the receptor and its ligand creates an intricate regulatory system that allows activation of the EGFR pathway at specific developmental time points^74^. In the lymph gland, cleavage of the spi ligand is activated by reactive oxygen species in the PSC during wasp parasitism, which then triggers the EGFR pathway to differentiate blood progenitors into lamellocytes^67, 68^. To prevent precocious differentiation, the EGFR pathway remains largely inactive during normal development in blood progenitors of intact lymph glands; however, the mechanisms underlying the developmental inactivation of EGFR are largely unidentified.

In this study, we found that Nos is expressed in blood progenitor cells in the lymph gland of *Drosophila* larvae and generates a gaseous ligand, nitric oxide (NO). NO is primarily utilized to S-nitrosylate target proteins in the core blood cell progenitors, which are in the most undifferentiated state. S-nitrosylation developmentally activates the Ire1–Xbp1-mediated UPR in the blood progenitors, thereby maintaining their multipotency downstream of NO. However, in the absence of NO or UPR activity, the core progenitor cells prematurely differentiate, or escape their undifferentiated state, into differentiating progenitors. Notably, we identified the EGFR as a key target of S-nitrosylation in the blood progenitors. S-nitrosylation of EGFR leads to it being trapped in the ER and inhibition of its membrane localization and function in blood progenitors. In summary, our findings suggest that NO is required for maintenance of the fate of blood progenitor cells through the developmentally programmed UPR induced by S-nitrosylation of target proteins, including EGFR, in the blood progenitor cells.

## Results

### Differential localization of EGFR in the lymph gland

By taking advantage of a CRISPR-GFP knock-in fly that specifically targets the C-terminus of the EGFR protein^75^, known as *EGFR-sfGFP*, we monitored the endogenous expression of EGFR in the lymph gland during larval development. Low levels of EGFR expression become visible during the mid-second instar (72 h after egg laying (AEL)), and later, this expression is increased in blood progenitor cells during the early-third instar (96 h AEL) (Figures 1A and 1B). By the late-third instar (120 h AEL), EGFR expression is maintained in the progenitor cells but at a lower level; it is less apparent compared with the early-third instar (Figure 1B). During the early-third instar, when EGFR expression is at its highest, it is mainly expressed in the core progenitors next to the dorsal vessel and co-localized with the core progenitor markers, *Tep4-Gal4*, col^Low^, or shg/E-cad (Figures 1C and 1E). Additionally, EGFR occupied an area a few cell diameters wider than that of col^Low^ or shg-positive core progenitor cells (Figures 1D and 1E) and is co-localized with the earliest differentiating blood cell marker, Pxn (Figure 1F), or an intermediate progenitor marker, *Nplp2*, at the distal margin (Figure 1G). Unlike progenitor cell markers, NimC1, a marker for mature plasmatocytes, and EGFR are mutually exclusive (Figures 1B and 1G), suggesting that EGFR is specifically expressed in lymph gland blood progenitor cells. In addition to its expression in the progenitors, high levels of EGFR are observed in the PSC (Figure 1D), accompanied by high levels of col, and in the posterior lobes, regardless of developmental stage (Figure S1A).

**Figure 1.**
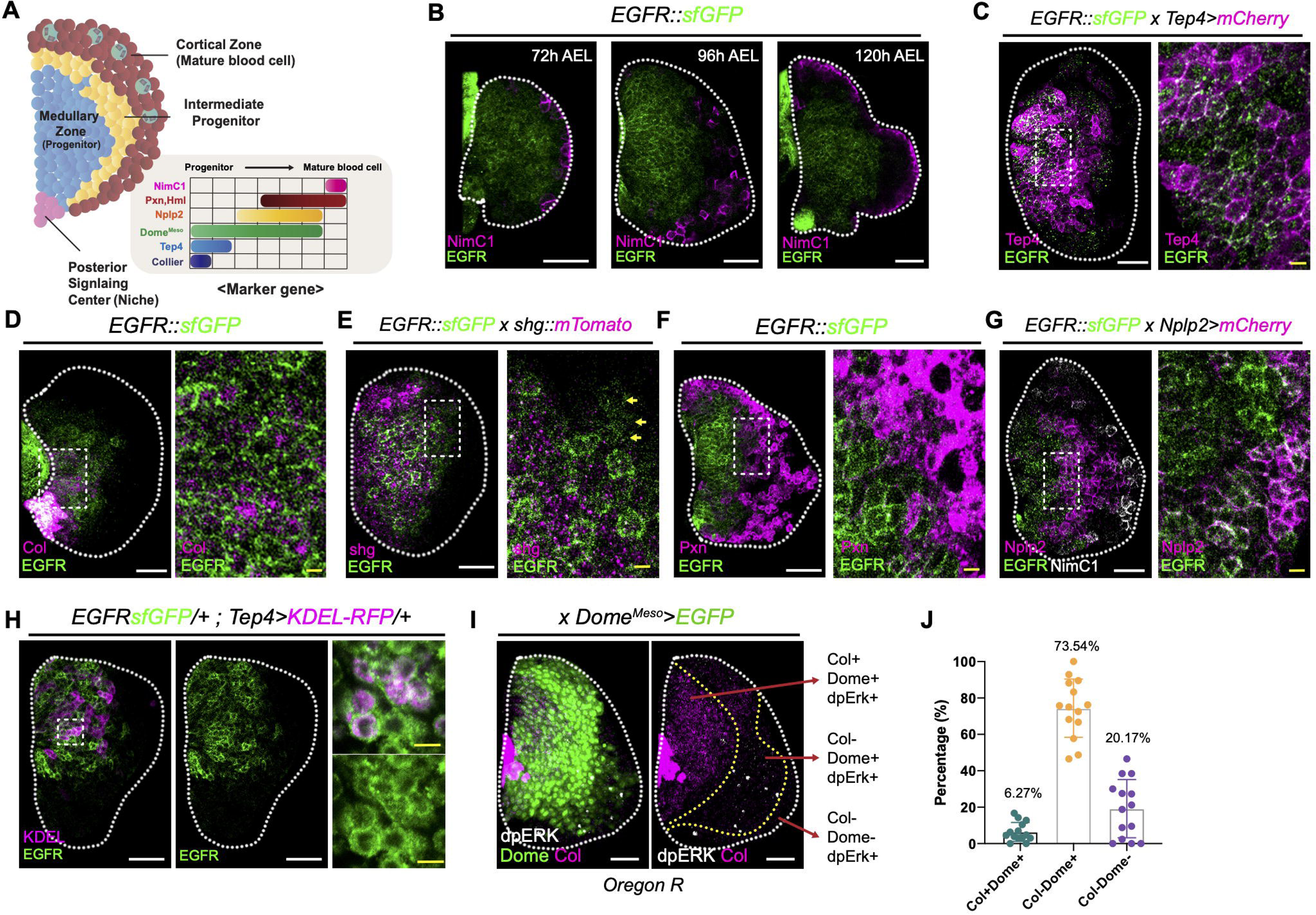
EGFR is expressed in the lymph gland blood progenitor cells. A. A schematic diagram of the *Drosophila* lymph gland (left). Progenitors are localized in the medial side of the lymph gland, comprising the medullary zone (blue). Differentiated mature blood cells place distal to the progenitors to form the cortical zone (red). Differentiating progenitor cells, called intermediate progenitors (yellow), are localized between the medullary and cortical zones. The posterior signaling center (PSC) is located in the posterior tip of the lymph gland. The gray box shows representative marker genes of lymph gland blood cells according to their degree of differentiation. Low-level collier (col^Low^) indicates the most primitive progenitor cells and *Tep4* overlaps with the col expression. *Dome^Meso^* is a pan-progenitor marker. *Nplp2* is a marker for intermediate progenitors. *Hml* or Pxn indicates blood cells in a differentiating state. **B-G.** EGFR protein expression in the lymph gland. Changes in the pattern of *EGFR-sfGFP* during development (*EGFR-sfGFP*) are indicated with a mature blood cell marker, NimC1 (magenta). EGFR (green) is expressed from 72 hours after egg laying (h AEL) and does not co-localize with NimC1 (magenta) **(B)**. EGFR (green) co-localizes with *Tep4* (magenta) (*EGFR-sfGFP, Tep4-Gal4 UAS-mCherry*) **(C)**. EGFR (green) is co-expressed with col (magenta) **(D)**. shotgun (magenta) overlaps with EGFR (green), but is expressed in a few cell diameters wider ranges (yellow arrow) than EGFR (*EGFRsfGFP, shg::mTomato*) **(E)**. A subset of EGFR-expressing cells (green) in their distal region co-localize with Pxn^+^ differentiating blood cells (magenta) **(F)**. EGFR (green) is partially co-localized with *Nplp2*^+^ intermediate progenitors (magenta) (*EGFR-sfGFP, Nplp2-Gal4 UAS-mCherry*). NimC1(white) **(G)**. **H.** In the blood progenitors at 120 h AEL, the majority of EGFR (green) is co-localized with an the ER marker, KDEL (magenta) (*EGFR-sfGFP Tep4-Gal4 UAS-KDEL-RFP*). **I-J.** Expression of dpErk in the lymph gland. Approximately 73% of dpERK-expressing cells are observed in col-negative, *Dome^Meso^*-positive progenitors **(I)**. Quantification of dpERK-positive cells according to progenitor markers **(J)**. *Dome^Meso^* (green), col (magenta), dpErk (white). Red arrows indicate markers and corresponding area. Magnified images of the insets in (C), (D), (E), (F), (G), and (H) are shown on the right of each panel. White scale bar, 30 μm. yellow scale bar, 3 μm. White dotted lines demarcate one lymph gland lobe.

By examining EGFR expression more closely, we noticed that most EGFR in blood progenitors is localized in the cytoplasm or in puncta, rather than on the cell membrane where it normally functions, during the mid-second or late-third instars (Figures 1B and 1E). Some of the core blood progenitors exhibit strong EGFR expression on the cell membrane during the early-third instar, but this pattern is attenuated in later stages (Figure 1B). Given that transmembrane receptor proteins are modified and transported to the membrane via the endoplasmic reticulum (ER) and Golgi^76–78^, we hypothesized that EGFR localizes to the ER in blood progenitors. Co-expression of *EGFR-sfGFP* with an ER indicator, *KDEL-RFP* (*Tep4-Gal4 UAS-KDEL-RFP, EGFR-sfGFP*), shows that progenitor cells have a high density of ER, and that most EGFR expression co-localizes with the ER marker (Figure 1H). In contrast, the PSC and other tissues, including the salivary gland and fat body, display strong EGFR expression on the cell membrane with minimal cytoplasmic expression (Figure S1B), implying that the subcellular localization of EGFR is modulated during progenitor development, possibly by distinct regulatory mechanisms.

Previous studies have shown that the EGFR signaling activity, indicated by the expression of phosphorylated ERK (dpERK), is largely attenuated in the lymph gland under conventional conditions^68^. Additionally, a heteroallelic combination of EGFR mutants does not interfere with lymph gland development^67^. Consistent with these findings, we confirmed that col^Low^-expressing core progenitors (col^Low+^*Dome^Meso^*^+^) are devoid of dpERK expression, whereas a few dpERK-expressing cells are found in differentiating progenitors (col^Low-^Dome^+^) or differentiated blood cells in the CZ (col^Low-^Dome^-^) (Figures 1I and 1J). Progenitor-specific expression of a dominant-negative form of EGFR (*Tep4-Gal4 UAS-EGFR^DN^*) does not alter lymph gland development (Figures S1C and S1D). However, expression of a constitutively active form of EGFR (*Tep4-Gal4 UAS-EGFR^Act^*) in the core progenitors dramatically induces differentiation of mature blood cells with a concomitant reduction in progenitors (Figures S1C and S1D). Together, these results suggest that EGFR is not exclusively localized in the membrane, and that the activity of the EGFR pathway is inactive in blood progenitors during their maintenance.

### Nitric oxide synthase is expressed in blood progenitors

Given the variations in EGFR localization and its attenuated signaling activity, we hypothesized that EGFR protein localization is altered in blood progenitors to sustain progenitor cell fates, enabling timely differentiation.

Nitric oxide (NO) is a highly diffusible messenger molecule that crosses cell membranes, but it also can locally regulate specific cellular processes, including the speed of protein transport from the ER to the cell membrane via protein modification^79^. In the lymph gland, NO has been implicated in myeloid blood cell development through a non-canonical Notch–Hifα interaction^80, 81^. However, it is unclear whether NO controls progenitor cell fate in a cell-autonomous manner and how it might accomplish this. Thus, we asked if NO controls blood progenitor fate by modifying protein targets. Using an antibody against *Drosophila* Nos, anti-Nos^82^, we analyzed the expression of Nos in the lymph gland. The Nos enzyme is enriched in blood progenitors expressing a pan-progenitor marker, *Dome^Meso^*(*Dome^Meso^-Gal4 UAS-EGFP*) (Figure 2A), which partially overlaps with the inner boundary of differentiating blood cells expressing *Hml* (*Hml*^Δ^*-Gal4 UAS-EGFP*) (Figure S2A). Nos protein expression is first detected at low levels from the mid-second instar at 72 h AEL when progenitor cells begin to differentiate^80^ (Figure S2B). Nos is gradually amplified until the mid-third instar at 96 h AEL and restricted to progenitor cells by the late-third instar at 120 h AEL, except for a few crystal cells in the CZ, concurrent with the establishment of the MZ and CZ (Figure S2B). We validated the specificity of the anti-Nos antibody with a heteroallelic combination of *Nos* mutants (*Nos*^Δ*all*^/*Nos*^Δ*15*^) and observed that Nos expression is eliminated in the mutant genetic background (Figure S2C).

**Figure 2.**
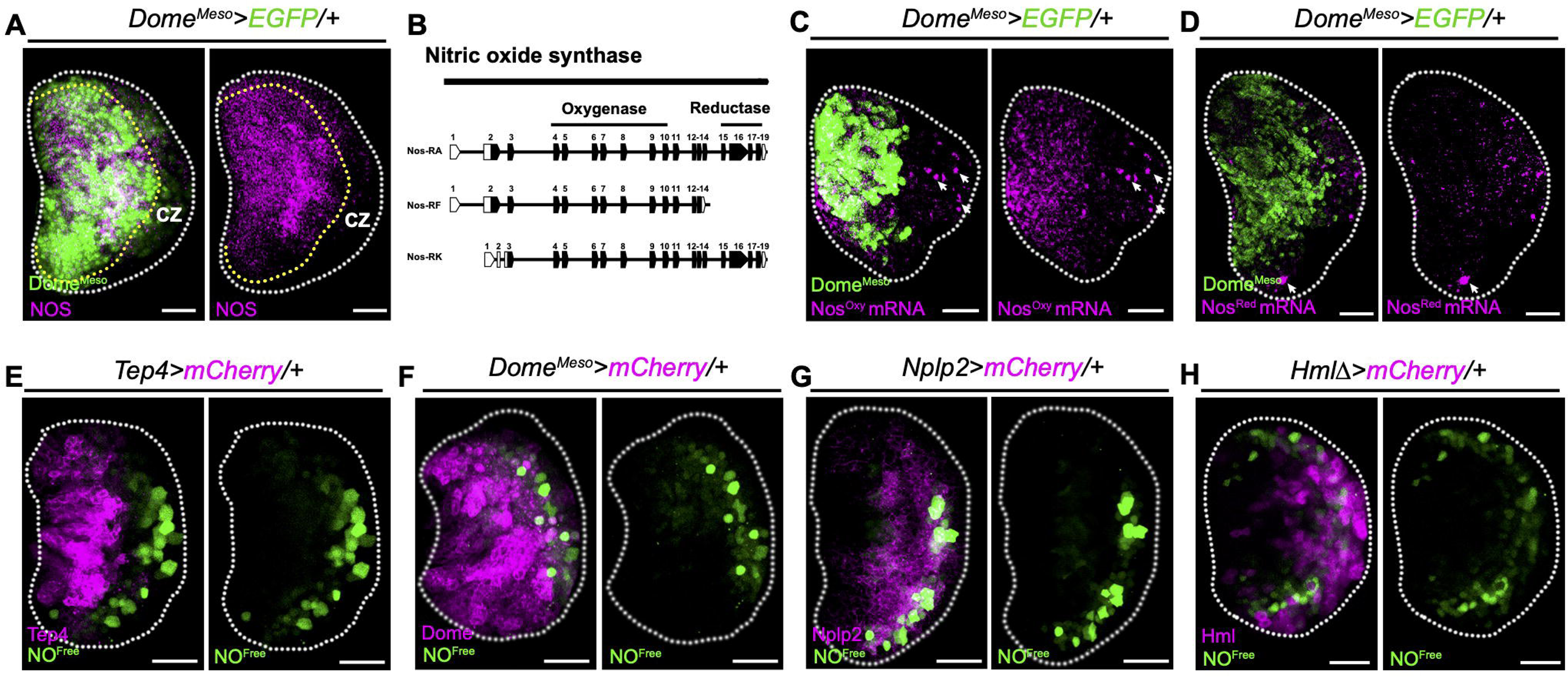
Nos and the free-radical NO are spatially segregated in lymph gland progenitors. **A.** Nos is expressed in blood progenitors. Nos (magenta) is expressed in *Dome^Meso^-*positive progenitors (*Dome^Meso^-Gal4 UAS-EGFP*). Yellow dotted line demarcates lymph gland progenitor cells. CZ indicates the cortical zone. **B.** Isoforms of *Nos* mRNA in *Drosophila melanogaster*. *Nos-RA* and *Nos-RK* encode both oxygenase (exons 4-10) and reductase (exons 15-18) domains, but *Nos-RF* lacks the reductase domain. Coding and untranslated regions of the mRNA are indicated by black and white boxes, respectively. **C-D.** Visualization of *Nos* mRNA by SABER-FISH. Oxygenase domain of *Nos* mRNA (*Nos^Oxy^*) (magenta) is expressed in *Dome^Meso^*-positive progenitors (green) or in crystal cells (white arrow) (*Dome^Meso^-Gal4 UAS-EGFP*) **(C)**. *Nos* reductase domain (*Nos^Red^*) (magenta) is absent in *Dome^Meso^*-positive progenitors (green) but is found in crystal cells (white arrow) (*Dome^Meso^-Gal4 UAS-EGFP*) **(D)**. **E-H.** Visualization of free-radical NO by DAF-FM diacetate (NO^Free^; green) with a progenitor cell marker, *Tep4* (magenta) (*Tep4-Gal4 UAS-mCherry*) **(E)**, a pan-progenitor marker, *Dome^Meso^* (magenta) (*Dome^Meso^-Gal4 UAS-mCherry*) **(F)**, an intermediate progenitor marker, *Nplp2* (magenta) (*Nplp2-Gal4 UAS-mCherry*) **(G)**, or a mature blood cell marker, *Hml* (magenta) (*Hml^Δ^-Gal4 UAS-mCherry*) **(H)**. White scale bar, 30 μm. White dotted line demarcates the lymph gland.

Although *Drosophila* encodes one *Nos* gene, three different isoforms—RA, RF, and RK—are annotated (FlyBase Release FB2019_96; http://flybase.org) (Figure 2B). Nos enzymes require two functional domains, a reductase and an oxygenase, which supply electrons and catalyze NO generation, respectively^83^. While the translated forms of *Nos-RA* and *Nos-RK* isoforms contain both domains, *Nos-RF* encodes a protein that lacks the C-terminal reductase domain essential for transferring electrons^84, 85^. To examine isoform-specific expressions of *Nos* mRNA in the lymph gland, we used SABER-FISH^86^ to differentially mark the N-terminal oxygenase and C-terminal reductase domains of *Nos* in the lymph gland. Interestingly, the *Nos* oxygenase domain (*Nos^Oxy^*) is readily detected in *Dome^Meso+^* progenitors (Figure 2C). However, we found that progenitors are devoid of *Nos* reductase domain (*Nos^Red^*) transcripts (Figure 2D). Only a subset of mature blood cells, including crystal cells, display both *Nos^Oxy^* and *Nos^Red^* (Figures 2C and 2D; Fig S2D). Thus, we conclude that blood progenitors in the lymph gland exhibit an isoform-specific expression of *Nos-RF* (*Nos^Oxy^*) during their development.

To identify if Nos^Oxy^ is functional and can generate NO in the lymph gland, we probed for the free radical form of NO using a live cell dye, DAF-FM-diacetate^87^. Notably, core progenitors, marked by *Tep4-Gal4 UAS-mCherry*, do not express free NO, despite their expression of Nos (Figure 2E). In contrast, a marker for the entire progenitor population, *Dome^Meso^* (*Dome^Meso^-Gal4 UAS-mCherry*), shows partial overlap with free-radical NO in a region distal to the core progenitors and those close to the edge of the differentiating progenitor population (Figure 2F). In addition, *Nplp2* (*Nplp2-Gal4 UAS-mCherry*), an IZ marker, or one that is characteristic of a differentiating blood cell marker, *Hml* (*Hml^Δ^-Gal4 UAS-mCherry*), shows evidence for the presence of free-radical NO (Figures 2G and 2H). Consistent with detection of the Nos protein, lymph glands from heteroallelic combinations of *Nos* mutants (*Nos^Δall^/Nos^Δ15^* or *Nos^1^/Nos^Δ15^*) lack free NO (Figure S2E). Together, these findings indicate that *Dome^Meso^*^+^ progenitors express an oxygenase-specific isoform of Nos in the lymph gland; however, only a subset of differentiating progenitors in the IZ contain free-radical NO.

### Nitric oxide and intracellular calcium promote S-nitrosylation in blood progenitor cells

In addition to its well-known function as a gaseous ligand, NO covalently modifies thiol groups of cysteine residues to produce S-nitrosylated cysteines (Cys-NOs) by a process called S-nitrosylation (S-NO)^15, 17^. We visualized Cys-NO-conjugated proteins in the lymph gland using an antibody against S-NO. Notably, S-NO-positive cells are specifically co-localized with *Tep4*^+^ in core progenitors but are excluded from intermediate progenitor cells (Figure 3A and 3B). During lymph gland development, low levels of S-NO begin in progenitors at 72 h AEL, which is gradually elevated by 96 h AEL but reduced again at 120 h AEL (Figure 3C). We validated the expression of the S-NO protein in the lymph gland by Western blotting using an S-NO-specific iodoTMT-labeling approach^15^. Consistent with expression detected by immunohistochemistry, the iodoTMT-labeled S-NO proteins are observed in lymph glands at all time points but attain their highest levels during the early-third instar (Figure S3A). *Nos*-dependent S-NO expression was confirmed in genetic assays using heteroallelic combinations of *Nos* mutants (*Nos^Δall^/Nos^Δ15^*) and an *Nos* RNA*i* clone (*Ay-Gal4 UAS-Nos RNAi*) (Figure 3D; Figure S3B). Based on these observations, we conclude that Nos generates NO in the lymph gland progenitors to facilitate S-NO of proteins in *Tep4*^+^ core blood progenitors.

**Figure 3.**
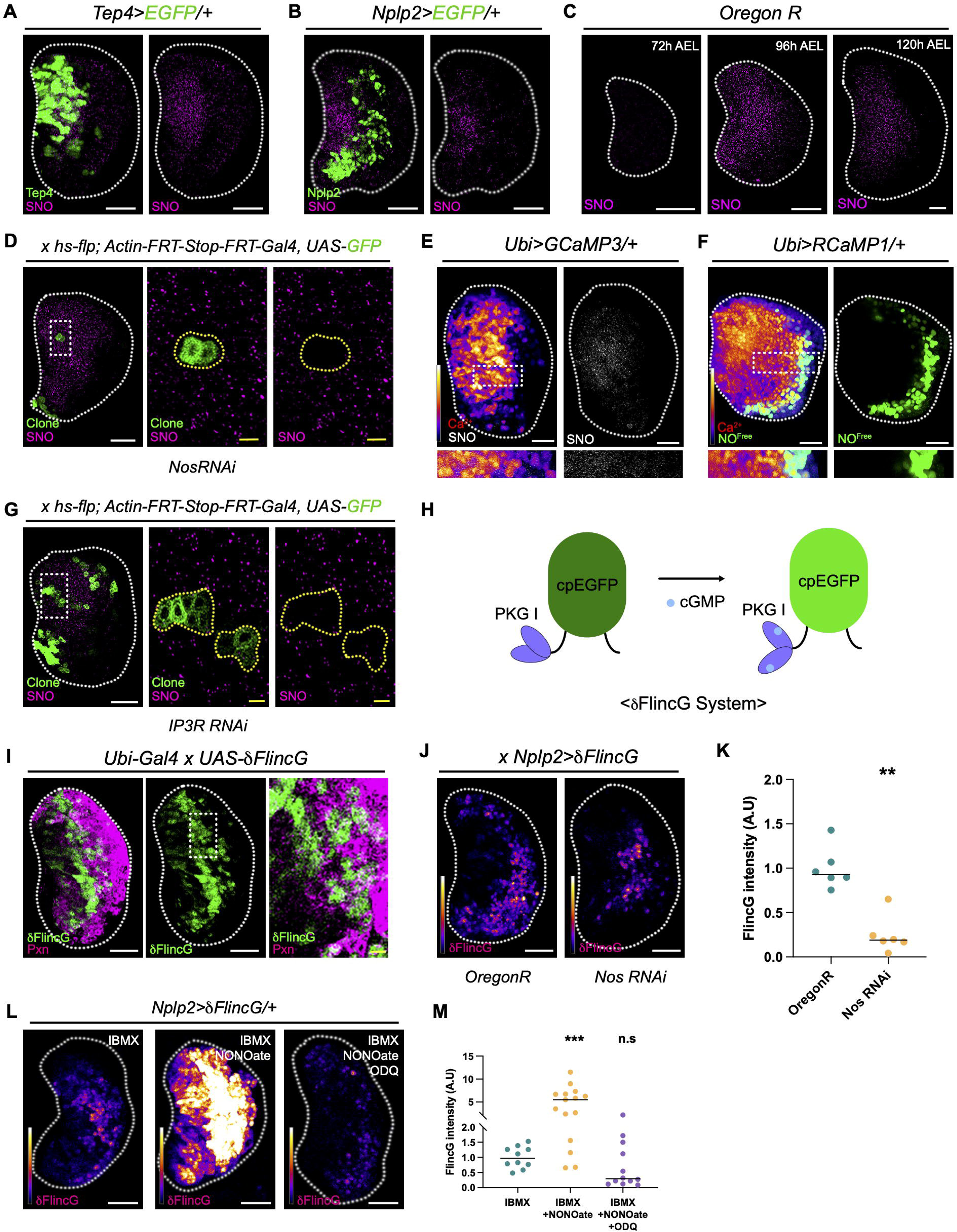
NO facilitates S-nitrosylation in core progenitors, but activates cGMP signaling in distal progenitors. A-B. Visualization of S-nitrosylation (SNO; magenta) with a core progenitor marker, *Tep4* (green) (*Tep4-Gal4 UAS-EGFP*) **(A)**, or with an intermediate progenitor marker, *Nplp2* (green) (*Nplp2-Gal4 UAS-EGFP*) **(B)**. The majority of SNO co-localize with *Tep4*^+^ progenitors, but not with *Nplp2^+^* progenitors. **C.** Visualization of S-nitrosylation (SNO; magenta) during the lymph gland development at 72, 96, or 120 h AEL. **D.** *Nos* RNAi clones (green; yellow dotted line) are devoid of S-nitrosylation (SNO; magenta) (*hs-flp, Actin-FRT-Stop-FRT-Gal4, UAS-GFP, UAS-NosRNAi)*. **E.** Visualization of S-nitrosylation (SNO; white) with a calcium level indicator, GCaMP3 (*Ubi-Gal4 UAS-GCaMP3*). Intracellular calcium levels are shown in the heatmap (left). S-nitrosylation correlates with high calcium expression in the lymph gland. A magnified image of an inset is shown below. **F.** Visualization of free-radical NO (NO^Free^; green) with a calcium level indicator, RCaMP1 (*Ubi-Gal4 UAS-RCaMP1*). Intracellular calcium levels are indicated in the heatmap (left). Free radical NO and high calcium expressions are mutually exclusive. A magnified image of an inset is shown below. **G.** *IP3R* RNAi expressing clones in blood progenitors (green; yellow dotted line) are devoid of S-nitrosylation (SNO; magenta) (*hs-flp, Actin-FRT-Stop-FRT-Gal4, UAS-GFP, IP3R RNAi*). **H.** Schematic illustration of the δFlincG sensor. cGMP binds to the PKG I domain of δFlincG and induces a conformational change of cpEGFP. δFlincG emits GFP fluorescence in the presence of cGMP. **I.** Visualization of intracellular cGMP using δFlincG (*Ubi-Gal4 UAS-δFlincG*). The majority of δFlincG signals (green) are displayed in the intermediate zone. A magnified image of an inset (middle) is shown on the right. Pxn (magenta). **J-K.** NO-dependent cGMP signaling in the intermediate progenitors. Reducing *Nos* expression in *Nplp2*^+^ differentiating progenitors decreases cGMP signaling in the lymph gland (*Nplp2-Gal4 UAS-δFlincG, UAS-NosRNAi*) **(J)**. Quantification of δFlincG intensity shown in (J) **(K)**. *Nplp2-Gal4 UAS-δFlincG* (n=6)*, Nplp2-Gal4 UAS-δFlincG, Nos RNAi* (n=6). **L-M.** NO-cGMP signaling is dependent on soluble guanylyl cyclase in the intermediate progenitors. An NO donor, NONOate, induces *δFlincG* signaling (middle), which is inhibited by a soluble guanylyl cyclase inhibitor, ODQ (right) **(L)**. Quantification of (L) **(M)**. White scale bar, 30 μm; yellow scale bar, 3 μm. White dotted lines demarcate one lymph gland lobe. Bars in graphs: the median. n.s.: not significant (p>0.01); \*\**p*<0.001 \*\*\**p*<0.0001.

Next, we asked how S-NO is specifically confined to the core progenitors and hypothesized that in addition to NO, another regulator may restrict S-NO specifically to the core progenitors. Past studies have highlighted the significance of calcium signaling in S-NO^24,88,89^. We therefore investigated whether high intracellular calcium levels in the progenitors, predicted by earlier studies^45, 46^, contribute to S-NO. Using the calcium indicators RCaMP1 and GCaMP3^90^, we compared the spatial distribution of progenitors expressing high calcium with those expressing free-radical NO or S-NO. We found that high calcium levels correlate well with S-NO (*Ubi-Gal4 UAS-GCaMP3*), whereas free-radical NO is detected in the outer boundaries of calcium-expressing cells and is absent from high-calcium areas (Figures 3E and 3F). To test if high calcium concentrations are required for S-NO, we generated clones expressing *IP3R* RNAi (*Ay-Gal4 UAS-IP3R RNAi*), which lowers intracellular calcium levels by blocking the transport of calcium from the ER to the cytosol^91^. In contrast to nearby wild-type clones (Figure 3G, GFP-negative), cells expressing *IP3R* RNAi have significantly less S-NO (Figure 3G, GFP-positive). Conversely, when *IP3R* expression is used to load high-calcium levels in the progenitor (*HHLT-Gal4 UAS-IP3R*), S-NO levels are not induced (Figure S3C), suggesting that intracellular calcium is required for S-NO in blood progenitors.

In neurons, free-radical NO acts as a signaling molecule and generates the second messenger cyclic GMP (cGMP) through activation of the NO-sensing soluble sGCs^12^. To investigate whether the sGC-mediated signaling cascade is triggered by free-radical NO in the IZ, we generated flies expressing a cGMP sensor, delta-FlincG^92^ (*UAS-δFincG*) (Figure 3H) and monitored cGMP levels in developing lymph glands. Despite its ubiquitous expression driven by *Ubi-Gal4*, the *δFincG* sensor is specifically expressed in Pxn^+^ differentiating blood cells at their proximal demarcation (*Ubi-Gal4 UAS-δFincG*) (Figure 3I; Figure S3D). Furthermore, we validated that RNAi-mediated knockdown of *Nos* in the IZ significantly reduces the intensity of *δFincG* sensor expression (*Nplp2-Gal4 UAS-δFincG UAS-Nos RNAi*) (Figure 3J and 3K), indicating the IZ-specific generation of cGMP depends on Nos. To establish whether cGMP is generated by stimulation of the sGC signaling cascade, we cultured the lymph gland expressing the *δFincG* sensor (*Nplp2-Gal4 UAS-δFincG*) ex vivo with chemicals modifying free-radical NO or the sGC pathway. As a control, we treated the lymph gland with a PDE inhibiter, IBMX, to stabilize the expression of the *δFincG* sensor in the ex vivo culture. Notably, the addition of an NO donor, NONOate, dramatically increases the number of *δFincG* sensor-expressing cells and the intensity of the *δFincG* signal (Figures 3L and 3M). Furthermore, simultaneous incubation of an sGC inhibitor, ODQ, with NONOate and IBMX suppresses the NONOate-mediated induction of *δFincG* activity (Figures 3L and 3M). Overall, these results establish that free-radical NO activates the sGC-mediated signaling pathway via cGMP in the IZ, while the core progenitors utilize NO to facilitate protein S-NO.

### S-nitrosylation attenuates the protein trafficking of EGFR in the blood progenitors

To identify the protein targets of S-NO, we used the iodoTMT-labeling technique to quantify S-NO in the lymph gland^93, 94^ (Figure 4A). In three biologically independent experiments using iodoTMT-labeling with tandem mass spectrometry, we found a variety of putative S-NO targets, including proteins involved in the cell cycle (Polo, sub, and mbt), RNA processing (AGO2, krimp), signaling cascades (e.g., Trol, Dl, and Uif), and immunity (Sr-CII, Tsf1) (Table S1). Of the identified proteins, EGFR was the most frequently found in all experiments, and cysteine 311 (C311) was the predominant target residue for S-NO (Figure 4B). C311 is located in the furine-like Cys-rich domain of the evolutionarily conserved extracellular region of EGFR essential for its ligand-dependent homodimerization^95^. Again, we confirmed the expression of S-nitrosylated EGFR in the lymph gland by immunoprecipitation of EGFR-sfGFP (Figure 4C). Moreover, we verified the in vivo subcellular co-localization of S-NO with EGFR-sfGFP in blood progenitor cells (Figure S4A). These results provide further proof that EGFR is S-nitrosylated in the blood progenitors of the lymph gland.

**Figure 4.**
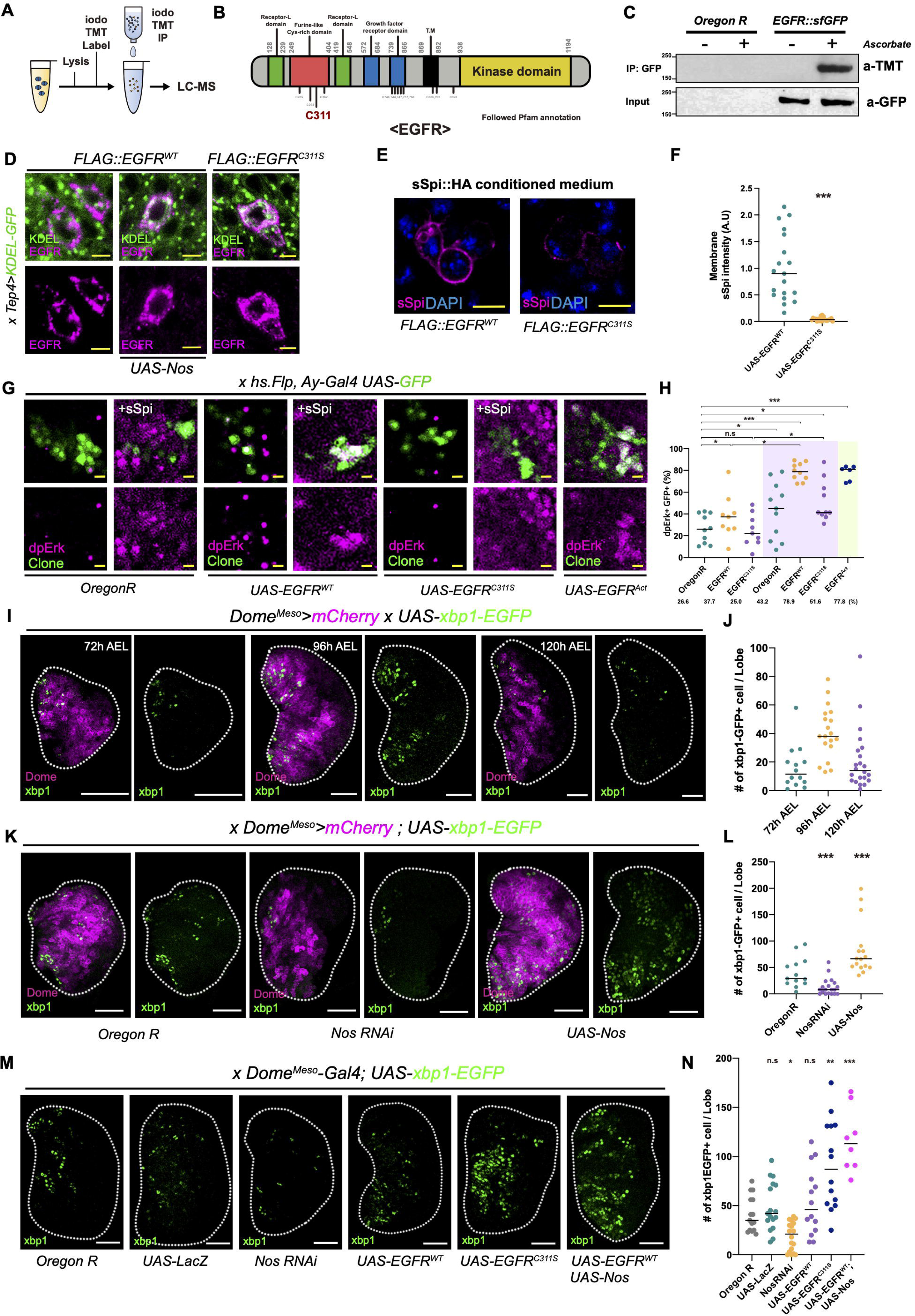
Ire1-dependent UPR pathway functions downstream of S-NO. A-B. Identification of EGFR S-nitrosylation in the lymph gland. Schematic representation of the TMT labeling procedure for LC-MS **(A)**. Immunoprecipitation of GFP-labeled EGFR protein (EGFR-sfGFP) with an anti-GFP antibody (input; bottom). S-nitrosylated EGFR is detected with an anti-TMT antibody (IP:GFP; top). Functional domains of the EGFR protein and localization of Cysteine 311 in the furine-like Cys-rich domain. **B.** Confirmation of S-nitrosylated EGFR in the lymph gland. Lymph glands expressing *EGFR-sfGFP* were TMT-labeled with or without ascorbate prior to immunoprecipitation **(B)**. *OreR* is a negative control. **D.** Localization of EGFR in the lymph gland. The majority of wild-type EGFR (EGFR^WT^) (magenta) is localized to the membrane (*Tep4-Gal4 UAS-KDEL-GFP UAS-FLAG::EGFR^WT^*) (left), whereas simultaneous overexpression of *Nos* (*Tep4-Gal4 UAS-KDEL-GFP UAS-FLAG::EGFR^WT^ UAS-Nos*) (middle) or the EGFR^C311S^ mutant (*Tep4-Gal4 UAS-KDEL-GFP UAS-FLAG::EGFR^C311S^*) (right) inhibits the membrane localization. In both cases, EGFR co-localizes with the ER marker, KDEL (green). **E-F.** EGFR^C311S^ mutants attenuate s.Spi ligand binding. s.Spi ligands (magenta) bind to wild-type EGFR (*FLAG::EGFR^WT^*) on the cell membrane of S2R^+^ cells (left). s.Spi ligands (magenta) do not bind to S2R^+^ cells expressing EGFR^C311S^ (*FLAG::EGFR^C311S^*) (right) **(E)**. DAPI (blue). Quantification of membrane-bound s.Spi intensity **(F)**. **G-H.** EGFR^C311S^ mutant protein does not induce dpErk in lymph gland clones. ∼ 26% of wild-type lymph gland clones (green; *hs-flp, Ay-Gal4 UAS-GFP*) show dpErk expression (magenta), which is increased by ex vivo culture with s.Spi conditioned medium (∼43%) (left). Lymph gland clones expressing EGFR^WT^ (*hs-flp, Ay-Gal4 UAS-GFP UAS-FLAG::EGFR^WT^*) increase dpERK expression up to ∼37%, which is further enhanced by ex vivo culture with s.Spi conditioned medium (∼78%) (middle, *UAS-EGFR^WT^*). Clones expressing EGFR^C311S^ (*hs-flp, Ay-Gal4 UAS-GFP UAS-FLAG::EGFR^C311S^*) do not show a significant increase by ex vivo culture with s.Spi (∼25% in normal, ∼51% in ex vivo culture with s.Spi) (middle, UAS-EGFR^C311S^). The constitutively active EGFR is used as a positive control (*hs-flp, Ay-Gal4 UAS-GFP UAS-EGFR^Act^*) **(G)**. Quantification of of dpERK^+^ GFP^+^ clones in panels (G) **(H)**. Purple shade in the graph indicates s.Spi treated samples. Green shade in the graph indicates positive controls. *Oregon R* (n=10), *EGFR^WT^* (n=9), *EGFR^C311S^* (n=9), *Oregon R* with s.Spi (n=11), *EGFR^WT^* with s.Spi (n=10), *EGFR^C311S^* with s.Spi (n=10), *EGFR^Act^* (n=6). **I-J.** Expression of *Xbp1-EGFP*, an ER stress marker, in *Dome^Meso^*-positive progenitors (*Dome^Meso^-Gal4 UAS-mCherry UAS-Xbp1-EGFP*) (magenta) during larval development at 72, 96, or 120 h AEL **(I)**. Quantification of Xbp1^+^ cells shown in panels (I) **(J).** 72 h (n=14), 96 h (n=19), 120 h (n=22). **K-L.** Nos-dependent expression of *Xbp1-EGFP* in *Dome^Meso^*-positive progenitors (*Dome^Meso^-Gal4 UAS-mCherry UAS-Xbp1-EGFP*) (magenta). Compared to controls (left), knockdown of *Nos* (*Dome^Meso^-Gal4 UAS-mCherry UAS-Xbp1-EGFP*, *UAS-NosRNAi*) (middle) or overexpression of *Nos* (*Dome^Meso^-Gal4 UAS-mCherry UAS-Xbp1-EGFP, UAS-Nos*) (right) reduces or increases the number of *Xbp1-EGFP*^+^ progenitors, respectively **(K)**. Quantification of the panels shown in (K) **(L)**. *Dome^Meso^-Gal4 UAS-mCherry UAS-Xbp1-EGFP* (n=13)*, Dome^Meso^-Gal4 UAS-mCherry UAS-Xbp1-EGFP, UAS-Nos RNAi* (n=19)*, Dome^Meso^-Gal4 UAS-mCherry UAS-Xbp1-EGFP, UAS-Nos* (n=16). **M-N.** S-nitrosylated EGFR induces *Xbp1-EGFP*^+^ cells in *Dome^Meso^*^+^ progenitors. Compared to wild-type (*Dome^Meso^-Gal4 UAS-mCherry UAS-Xbp1-EGFP)* or LacZ-expressing control lymph glands (*Dome^Meso^-Gal4 UAS-mCherry UAS-Xbp1-EGFP, UAS-LacZ*), overexpression of EGFR^WT^ (*Dome^Meso^-Gal4 UAS-mCherry UAS-Xbp1-EGFP, UAS-FLAG::EGFR^WT^*) does not alter the number of *Xbp1-EGFP*^+^ cells (green). Expression of EGFR^C311S^ (*Dome^Meso^-Gal4 UAS-mCherry UAS-Xbp1-EGFP, UAS-FLAG::EGFR^C311S^*) or overexpression of wild-type EGFR^WT^ with Nos (*Dome^Meso^-Gal4 UAS-mCherry UAS-Xbp1-EGFP, UAS-FLAG::EGFR^WT^ UAS-Nos*) significantly increases the number of *Xbp1-EGFP*^+^ cells (green). Expression of RNAi against *Nos* (*Dome^Meso^-Gal4 UAS-mCherry UAS-Xbp1-EGFP, UAS-NosRNAi*) decreases the number of *Xbp1-EGFP*^+^ cells **(M)**. Quantification of *Xbp1-EGFP*^+^ cells in the panels shown in (M) **(N)**. *Oregon R* (n=15), *UAS-LacZ* (n=17), *NosRNAi* (n=18), *EGFR^WT^*(n=14), *EGFR^C311S^* (n=14), *EGFR^WT^ UAS-Nos* (n=8). White scale bar, 30 μm; yellow scale bar, 3 μm. White dotted lines demarcate the lymph gland n.s.: not significant (p>0.01); \**p*<0.01; \*\**p*<0.001, \*\*\**p*<0.0001. Bars in graphs: the median.

S-NO is a post-translational modification that disrupts disulfide bonds critical for protein folding, potentially trapping target proteins in the ER^15, 17^. Based on our observations, we hypothesize that S-NO may modify the localization of the EGFR protein in blood progenitor cells. To test this, we overexpressed Nos in the core progenitors and analyzed the cellular localization of EGFR. The core progenitor-specific expression of Nos significantly inhibits the membrane localization of EGFR, instead retaining it in the cytoplasm, where most of it co-localizes with the ER marker, KDEL (*Tep4-Gal4 UAS-KDEL-GFP UAS-Nos UAS-FLAG-EGFR^WT^*) (Figure 4D). This phenomenon is more clearly observed in the giant cells of the salivary gland, where the majority of EGFR is found in the membrane in wild-type tissues (Figure S4B). To further confirm the role of S-NO in the localization of EGFR, we created a point mutation that mimics S-NO by replacing EGFR C311 with serine (EGFR^C311S^). Similar to the expression of Nos, expression of EGFR^C311S^ in core progenitors induces the ER retention of EGFR protein (*Tep4-Gal4 UAS-KDEL-GFP UAS-FLAG-EGFR^C311S^*) (Figure 4D). Again, this phenotype is recapitulated in the salivary gland (Figure S4B). These results indicate that S-NO of EGFR blocks its translocation to the plasma membrane and causes its accumulation in the ER.

Next, we analyzed the functional relevance of S-nitrosylated EGFR in ligand-binding and its signaling cascade by expressing either *EGFR^WT^* or *EGFR^C311S^* in *Drosophila* embryonic hemocyte-derived S2R^+^ cells (Figure S4C). After culturing the cells with an HA-tagged secreted form of the spi protein (s-spi), a potent ligand for *Drosophila* EGFR^96, 97^, s-spi strongly binds to the surface of EGFR^WT^-expressing S2R^+^ cells, but expression of EGFR^C311S^ significantly disrupts the localization of s-spi to the membrane (Figures 4E and 4F). Similarly, in EGFR^WT^-expressing S2R^+^ cells, dpERK is observed after s-spi treatment, and this too is inhibited upon EGFR^C311S^ expression (Figures S4D and S4E). This demonstrates that the S-NO mimic blocks signaling by EGFR when targeted to the membrane. Finally, to confirm this role of S-nitrosylated EGFR in the lymph gland, we generated *Ay-Gal4* clones, which produces random clones by heat shock-activated Flippase^98^, and cultured the lymph gland with s-spi ex vivo to assess EGFR activity using dpERK as a marker. Most EGFR^WT^-expressing clones in dissected lymph glands not treated with spi do not show dpERK expression, likely suggesting that the source of s-spi is normally from the hemolymph surrounding the lymph gland (*hs-FLP Ay-Gal4 UAS-GFP UAS-EGFR^WT^*) (Figures 4G and 4H). Consistent with this hypothesis, clones of *EGFR^WT^* exhibit dpERK expression when the medium is treated with s-spi protein (Figures 4G and 4H). Also, as positive controls, clones of *EGFR^Act^*, a constitutive, ligand-independent form of EGFR, show clear signs of dpERK induction, even in the absence of exogenously provided s-spi (*hs-FLP Ay-Gal4 UAS-GFP* or *hs-FLP Ay-Gal4 UAS-GFP UAS-EGFR^Act^*) (Figures 4G and 4H). In contrast, clones expressing *EGFR^C311S^* do not exhibit dpERK induction, even with s-spi treatment (*hs-FLP Ay-Gal4 UAS-GFP UAS-EGFR^C311S^*) (Figures 4G and 4H). Overall, these results demonstrate that S-nitrosylated EGFR inhibits ligand-binding and downstream activity when present in the cell membrane before resolving the modification.

### S-nitrosylation activates the unfolded protein response in blood progenitors

Given that S-NO alters protein localization and proper trafficking in progenitors, we examined whether the core progenitors experience the UPR during development. UPR is well-conserved across species and consists of three parallel pathways that aim to resolve ER stress: 1) Ire1–Xbp1, 2) PERK/Atf4, and 3) Atf6 pathways^26, 99^. To measure the activity of the Ire1–Xbp1 pathway, we used the *Xbp1-EGFP* reporter, which monitors the Ire1-dependent alternative splicing of *Xbp1*mRNA^22^. Despite the ubiquitous expression of *UAS-Xbp1-EGFP* throughout the entire lymph gland (*Ubi-Gal4 UAS-Xbp1-EGFP*), most *Xbp1-EGFP* expression is observed in core progenitors expressing col^Low^, while only a small subset is found in Pxn^+^ differentiating blood cells (Figures S5A and S5B). Developmentally, the expression of *Xbp1-EGFP* is initiated by the late-second instar at 72 h AEL, amplified by 96 h AEL, and diminished by 120 h AEL, reminiscent of the pattern observed with anti-S-NO (*Dome^Meso^-Gal4 UAS-Xbp1-EGFP*) (Figures 4I and 4J; Figure 3C; Figure S3A). Next, we examined the expression of Atf4 (crc in *Drosophila*) as a readout for the PERK pathway using an antibody against Atf4^25^. We found that Atf4 is expressed in *Dome^Meso^*-negative mature blood cells but not in the blood progenitors (Figures S5C). We validated the specificity of the anti-Atf4 antibody in the lymph gland by *Atf4* RNAi (*HHLT-Gal4 UAS-Atf4 RNAi*) and found that *Atf4* RNAi significantly reduces Atf4 expression (Figure S5D). Furthermore, we confirmed an increase in Atf4 expression in differentiated blood cells after being fed with an UPR inducer, dithiothreitol (DTT) (Figure S5E), indicating that the Atf4-mediated UPR pathway is functional primarily in differentiated blood cells. Finally, we measured the activity of the Atf6 pathway using *Atf6-GFP*^100^. Similar to the results with Atf4, DTT-mediated UPR activation successfully enhances the expression of Atf6-GFP in other tissues, such as the salivary gland; however, blood cells in the lymph gland do not express Atf6, even in the presence of DTT (Figure S5F), suggesting that the Atf6 pathway is inactive in the lymph gland.

We then investigated whether the Ire1–Xbp1-mediated UPR pathway is dependent upon NO. When *Nos* is downregulated in *Dome^Meso^*-expressing progenitors, the number of *Xbp1-EGFP*-expressing progenitors is significantly reduced in the lymph gland (*Dome^Meso^-Gal4 UAS-mCherry UAS-Xbp1-EGFP UAS-NosRNAi*) (Figures 4K and 4L). We also found that overexpression of *Nos* considerably elevates the level of *Xbp1-EGFP*. In addition, changes in the expression of Ire1–Xbp1 target genes were validated using qRT-PCR in the abovementioned genetic backgrounds (Figure S5G)^28^. Next, since EGFR is one of the targets of S-NO retained in the ER, we probed for a possible role of EGFR S-NO in inducing the Xbp1-dependent UPR response observed in blood progenitors. We found that expression of wild-type EGFR alone does not activate splicing of *Xbp1* in progenitors (*Dome^Meso^-Gal4 UAS-mCherry UAS-Xbp1-EGFP UAS-EGFR^WT^*), as compared with the control (*Dome^Meso^-Gal4 UAS-mCherry UAS-Xbp1-EGFP UAS-LacZ*) (Figures 4M and 4N). However, coexpression of *EGFR^WT^* and *Nos* leads to a significant increase in the number of *Xbp1-EGFP*-positive cells (*Dome^Meso^-Gal4 UAS-mCherry UAS-Xbp1-EGFP UAS-EGFR^WT^ UAS-Nos*) (Figures 4M and 4N). Similarly, expressing *EGFR^C311S^* in the blood progenitors also results in enhancement of the Ire1–Xbp1 pathway (*Dome^Meso^-Gal4 UAS-mCherry UAS-Xbp1-EGFP UAS-EGFR^C311S^*) (Figures 4M and 4N). Furthermore, we verified that downstream target genes of the UPR pathway are elevated by *EGFR^C311S^*expression but not by *EGFR^WT^*alone (Figure S5H). Taken together, these results establish that S-NO perturbs target protein localization, including EGFR, and activates the UPR pathway through the Ire1–Xbp1 signaling cascade as part of the core progenitor development.

### The S-nitrosylation/unfolded protein response axis maintains the blood progenitors

#### 1) S-nitrosylation is required for blood progenitor maintenance

We explored the function of *Nos* in blood progenitor maintenance by analyzing the proportions of progenitors or mature blood cells in *Nos*-mutant backgrounds. Under normal conditions, lymph glands undergo a stereotypical developmental pattern that results in an approximately 60-to-40 ratio of progenitors (marked with *Dome^Meso^*) to differentiated blood cells (marked with Pxn), respectively, at the late-third instar (Figures 5A and 5B). However, when *Nos* function is disrupted utilizing either of two independent heteroallelic combinations (*Nos*^Δ*all*^/*Nos*^Δ*15*^ or *Nos^1^*/*Nos*^Δ*15*^), the proportion of Pxn^+^ differentiating blood cells increases, and the expression of col^Low^-expressing core progenitors is significantly decreased (Figures 5A–5D). Although Pxn^+^ blood cells increase in number, the proportion of NimC1^+^ mature plasmatocytes is decreased (Figures S6A and S6B). These results suggest that Nos is important for preventing differentiation of the core progenitors. To investigate the role of S-NO in blood progenitors, we used RNAi to knock down the *Nos* reductase (*Nos^Red^*) or oxygenase domain (*Nos^Oxy^*) in *Dome^Meso^*^+^ progenitors (*Dome^Meso^-Gal4 UAS-Nos^Red^ RNAi* or *Dome^Meso^-Gal4 UAS-Nos^Oxy^* RNAi). Consistent with *Nos* mutants that enhance Pxn^+^ blood cells (Figures 5A and 5B), knocking down *Nos^Oxy^*, but not *Nos^Red^*, leads to an expansion of both Pxn^+^ differentiating blood cells and *Dome^Meso^*^+^ progenitors (Figures 5E–5G). *Dome^Meso^*^+^ progenitors at their distal margin give rise to cells that express both *Dome^Meso^* and *Hml*/Pxn, defined as the IZ undergoing differentiation^35^. Given an increase in both *Dome^Meso^* and Pxn in the *Nos^Oxy^* RNAi background (*Dome^Meso^-Gal4 UAS-Nos^Oxy^ RNAi*), we analyzed proportions of *Dome^Meso^*^+^ and Pxn^+^ double-positive blood cells and found that these double-positive cells, presumably forming an IZ, are consequently expanded in this genetic background (Figures S6C and S6D). In addition, expression of *Nos^Oxy^* RNAi in *Dome^Meso^*^+^ progenitors dramatically reduces the expression of col^Low+^ core progenitor cells (Figures 5H and 5I). However, a *Dome^Meso^*^+^ progenitor-specific knockdown of *Nos^Oxy^* or *Nos^Red^*does not affect the differentiation of NimC1^+^ plasmatocytes, which is reminiscent of phenotypes observed in *Nos* mutants (Figures S6E and S6F). Since S-NO is restricted to the core progenitors, we combined a driver, *Tep4-Gal4,* which is restricted to the core progenitors^53,54,101^, with *Nos^Oxy^*or *Nos^Red^* and found that *Tep4-Gal4* recapitulates the phenotypes observed with *Dome^Meso^-Gal4* (*Tep4-Gal4 UAS-Nos^Red^ RNAi* or *Tep4-Gal4 UAS-Nos^Oxy^ RNAi*) (Figures 6J–6O; Figures S6G and S6H). As expected, expression of RNAi against *Nos^Oxy^* driven by an intermediate progenitor-specific *Nplp2-Gal4* (*Nplp2-Gal4 UAS-Nos^Oxy^RNAi*) has no effect on the differentiation of Pxn^+^ differentiating blood cell phenotypes (Figures S6I and S6J).

**Figure 5.**
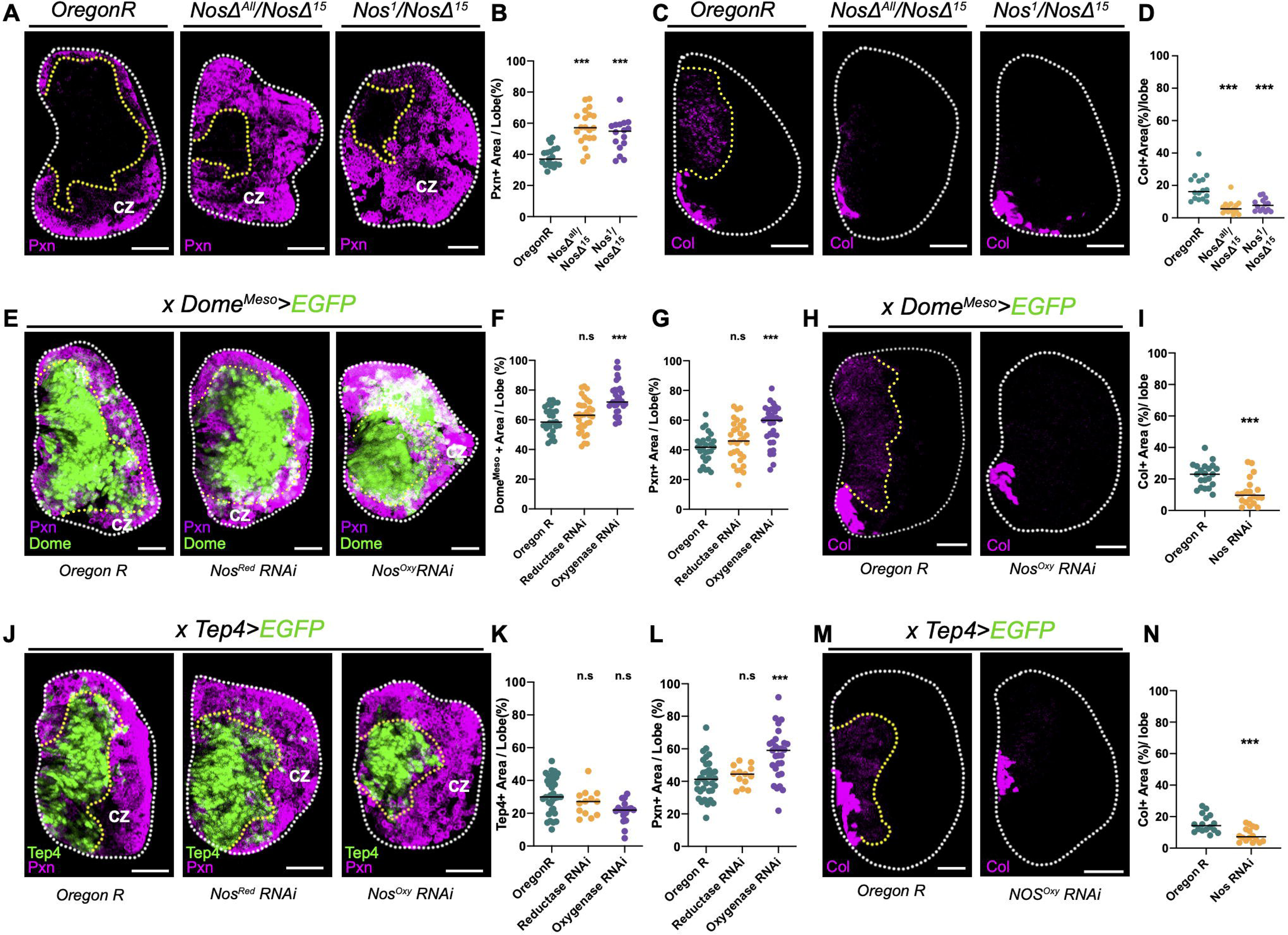
Nos is required for the maintenance of lymph gland progenitors. A-B. Differentiation of *Nos* mutants in the lymph gland. Compared to wild-type controls, hetero-allelic combinations of *Nos* mutants (*NosΔ^All^/NosΔ^15^* or *Nos^1^/NosΔ^15^*) display an increased proportion of Pxn^+^ differentiating blood cells (magenta) **(A)**. Quantification of Pxn^+^ area per one lymph gland lobe shown in panels (A) **(B)**. Yellow dotted lines indicate Pxn-negative progenitors. CZ indicates the cortical zone. *Oregon R* (n=18), *NosΔ^All^/NosΔ^15^*(n=20)*, Nos^1^/NosΔ^15^* (n=16). **C-D**. Expression of low-level col in *Nos* mutant lymph glands. Hetero-allelic combinations of *Nos* mutants (*NosΔ^All^/NosΔ^15^* or *Nos^1^/NosΔ^15^*) reduce the expression of low-level col, a core progenitor marker (magenta) **(C)**. Yellow dotted lines indicate low-level col^+^ progenitors. High col expression in the posterior side of the lymph gland indicates the posterior signaling center (PSC). Quantification of col^+^ area in panels (C) **(D)**. *Oregon R* (n=16), *NosΔ^All^/NosΔ^15^*(n=14)*, Nos^1^/NosΔ^15^* (n=14). **E-G**. Domain-specific knockdown of *Nos* in *Dome^Meso^*^+^ progenitors. Compared to wild-type lymph glands, knockdown of *Nos* oxygenase (*Nos^Oxy^*; BL28792) increases both *Dome^Meso^*^+^ progenitors (green) and Pxn^+^ differentiating blood cells (magenta) (*Dome^Meso^-Gal4 UAS-EGFP, Nos^Oxy^ RNAi*). Expression of an RNAi against *Nos* reductase (*Nos^Red^*; VDRC 27722) does not alter lymph gland development (*Dome^Meso^-Gal4 UAS-EGFP, UAS-Nos^Red^RNAi*). Yellow dotted lines demarcate the lymph gland progenitor cells. CZ indicates the cortical zone **(E)**. Quantification of *Dome^Meso+^* progenitors **(F)** or Pxn^+^ differentiating blood cells per one lymph gland lobe **(G)**. *Oregon R* (n=27), *Nos reductase RNAi* (n=32), *Nos oxygenase RNAi* (n=30). **H-I**. Progenitor-specific knockdown of *Nos^Oxy^* reduces low-level col expression (*Dome^Meso^-Gal4 UAS-EGFP, UAS-Nos^Oxy^RNAi*) **(H)**. Yellow dotted line indicates col^+^ progenitor cells. High col^+^ cells represent the posterior signaling center (PSC). Quantification of col^+^ cells in panels (H) **(I)**. *Oregon R* (n=21), *Nos oxygenase RNAi* (n=20). **J-L**. Domain-specific knockdown of *Nos* in *Tep4*^+^ progenitors. Compared to wild-type lymph glands, knockdown of *Nos oxygenase* (*Nos^Oxy^*; BL 28792) increases Pxn^+^ differentiating blood cells (magenta) (*Tep4-Gal4 UAS-EGFP, UAS-Nos^Oxy^RNAi*). Expression of RNAi against *Nos reductase* (*Nos^Red^*; VDRC 27722) does not modify lymph gland development (*Tep4-Gal4 UAS-EGFP, UAS-Nos^Red^RNAi*) **(J)**. Yellow dotted lines demarcate the lymph gland progenitor cells. CZ indicates the cortical zone. Quantification of *Tep4*^+^ progenitors **(K)** or Pxn^+^ differentiating blood cells **(L)**. *Tep4* marks the most undifferentiated progenitors, similar to col, different from a pan-progenitor marker, *Dome^Meso^*. *Oregon R* (n=32), *Nos reductase RNAi* (n=13), *Nos oxygenase RNAi* (n=29). **M-N.** *Tep4*^+^ progenitor-specific knockdown of *Nos^Oxy^*reduces the expression of col (*Tep4-Gal4 UAS-EGFP, UAS-Nos^Oxy^RNAi*) **(M)**. Yellow dotted lines indicate col^+^ progenitor cells. High col^+^ cells indicate the posterior signaling center (PSC). Quantification of on col^+^ cells in panels (M) **(N)**. *Oregon R* (n=21), *Nos oxygenase RNAi* (n=20). White scale bar, 30 μm. White dotted lines demarcate the lymph gland. n.s.: not significant (p>0.01); ***p<0.0001. Bars in graphs: the median.

**Figure 6.**
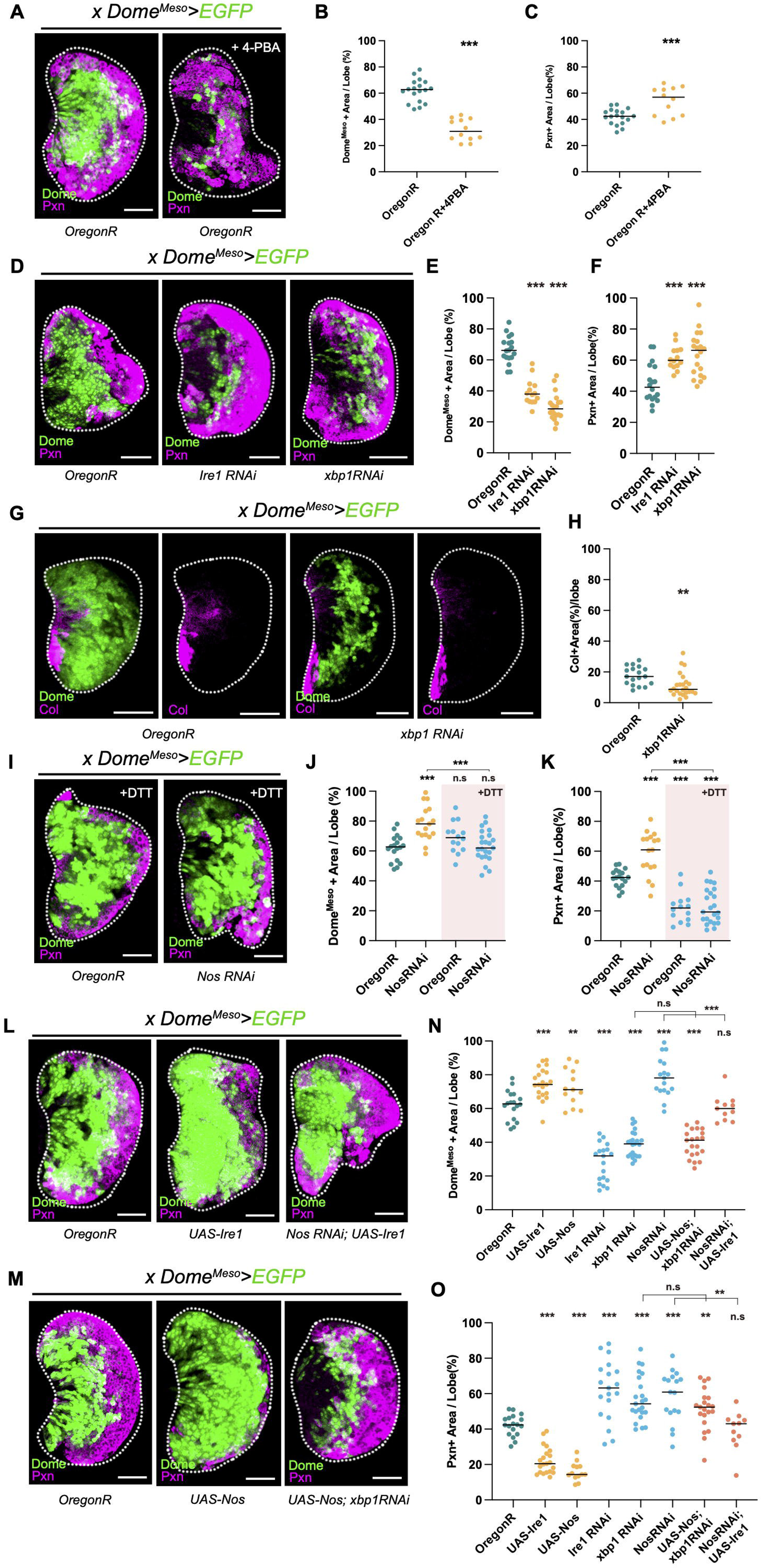
Ire1/Xbp1-dependent UPR functions downstream of S-NO. A-C. UPR is required for the progenitor maintenance in the lymph gland. Compared to wild-type controls, feeding 4-PBA, a chemical chaperone, decreases *Dome^Meso^*^+^ progenitors (green) and increases Pxn^+^ differentiating blood cells (magenta) **(A)**. Quantification of *Dome^Meso^*^+^ **(B)** or Pxn^+^ cells **(C)** per one lymph gland lobe after 4-PBA feeding. *Oregon R* (n=18), *Oregon R* with 4-PBA (n=11) **D-F.** Inhibition of the Ire1/Xbp1-dependent UPR pathway reduces the blood progenitors. Knockdown of *Ire1* (middle) or *Xbp1* (right) in the progenitor (*Dome^Meso^-Gal4 UAS-EGFP UAS-Ire1RNAi* or *Dome^Meso^-Gal4 UAS-EGFP UAS-Xbp1RNAi*) decreases *Dome^Meso^*^+^ progenitors (green) and increases Pxn^+^ differentiating blood cells (magenta) **(D)**. Quantification of *Dome^Meso^*^+^ progenitors **(E)** or Pxn^+^ differentiating blood cells **(F)** per one lymph gland lobe. *Oregon R* (n=18), *Ire1 RNAi* (n=14), *Xbp1 RNAi* (n=20). **G-H.** *Xbp1* is required for the maintenance of col^+^ core progenitors. Compared to wild type, *Dome^Meso^*^+^-specific knockdown of *Xbp1* decreases col^+^ core progenitor cells (magenta) (*Dome^Meso^-Gal4 UAS-EGFP UAS-Xbp1RNAi*) **(G)**. Quantification of col^+^ progenitors in (G) **(H).** *Oregon R* (n=18), *Xbp1 RNAi* (n=24). **I-K.** *Nos* RNAi-mediated progenitor differentiation is rescued by feeding DTT, a UPR inducer. Compared to wild-type lymph glands shown in (A), DTT feeding reduces Pxn^+^ differentiating blood cells (magenta) without altering the ratio of Dome^Meso+^ progenitors (green) (*Dome^Meso^-Gal4 UAS-EGFP*) (left). The increase in differentiating blood cells caused by *Nos* RNAi is recovered by DTT administration (*Dome^Meso^-Gal4 UAS-EGFP UAS-NosRNAi*) (right) **(I)**. Quantification of *Dome^Meso^*^+^ progenitors **(J)** or Pxn^+^ differentiating blood cells **(K).** *Oregon R* (n=18), *Oregon R* with DTT (n=14), *Nos RNAi* (n=17), *Nos RNAi* with DTT (n=23). Pink shade (+DTT) indicates DTT feeding. **L-O.** *Nos* RNAi-mediated progenitor differentiation is rescued by Ire1. Overexpression of *Ire1* alone in *Dome^Meso+^* progenitors inhibits Pxn^+^ blood cell (magenta) differentiation and expands *Dome^Meso+^* progenitors (green) (*Dome^Meso^-Gal4 UAS-EGFP UAS-Ire1*). Similarly, overexpression of *Ire1* with *Nos RNAi* in the progenitor rescues the *Nos* RNAi-driven increase in Pxn^+^ differentiation (*Dome^Meso^-Gal4 UAS-EGFP UAS-Ire1 UAS-NosRNAi*) **(L)**. Nos overexpression expands *Dome^Meso+^* progenitors (green) and inhibits Pxn^+^ differentiating blood cells (magenta) (*Dome^Meso^-Gal4 UAS-EGFP UAS-Nos*) but does not rescue the *Xbp1 RNAi* phenotype (*Dome^Meso^-Gal4 UAS-EGFP UAS-Nos UAS-Xbp1RNAi*) **(M)**. Quantification of *Dome^Meso^*^+^ area on **(N)** and Pxn^+^ area on **(O)**. *Oregon R* (n=21), *UAS-Ire1* (n=21), *UAS-Nos* (n=13), *UAS*-*Ire1 RNAi* (n=19), *UAS*-*Xbp1RNAi* (n=23), *UAS-Ire1 UAS-NosRNAi* (n=11)*, UAS-Nos UAS-Xbp1RNAi* (n=21). White scale bar, 30 μm; yellow scale bar, 3 μm. White dotted lines demarcate the lymph gland n.s: not significant (p>0.01); \**p*<0.01; \*\**p*<0.001, \*\*\**p*<0.0001. Bars in graphs: the median.

Concurrent with our findings that loss of intracellular calcium by *IP3R* RNAi inhibits S-NO expression (Figure 3G), modifying calcium levels in blood progenitors increases Pxn^+^ blood cell differentiation, consistent with our earlier findings^46^ (*Dome^Meso^-Gal4 IP3R RNAi*) (Figures S6K–S6M). From these results, we conclude that the expression of *Nos^Oxy^* in the core progenitors facilitates S-NO modifications, which is essential for the maintenance of core progenitor cells.

#### 2) Ire1–Xbp1-mediated UPR maintains blood progenitor cell fate

To investigate the significance of an active UPR in blood progenitor maintenance, we attenuated UPR activity by feeding the chemical chaperone 4-phenyl butyric acid (4-PBA) to larvae beginning at 72 h AEL until 120 h AEL, when S-NO expression is evident. Compared with controls, feeding 4-PBA significantly reduces *Dome^Meso^*^+^ progenitor cells and increases Pxn^+^ differentiating blood cells (*Dome^Meso^-Gal4 UAS-EGFP*) (Figures 6A–6C). Knockdown of *Ire1* or *Xbp1* in progenitors (*Dome^Meso^-Gal4 UAS-Ire1 RNAi* or *Dome^Meso^-Gal4 UAS-Xbp1 RNAi*) reproduces the differentiation phenotype induced by 4-PBA administration (Figures 6D–6F). In addition, inhibition of *Xbp1* in *Dome^Meso^*^+^ progenitors dramatically decreases col^Low^ expression (Figures 6G and 6H), while the differentiation of NimC1^+^ plasmatocytes remains unchanged (Figures S7A and S7B). These phenotypes are reminiscent of those observed with *Nos* RNAi (Figures S6E and S6F). To determine whether these phenotypes are specific to the core progenitors, we used the *Tep4-Gal4* driver and observed virtually identical phenotypes (Figures S7C-S7E). This confirms that the Ire1–Xbp1-mediated UPR is responsible for maintaining the core progenitors. Consistent with the lack of *Atf4* or *Atf6* expression in blood progenitors, reducing the expression of *PERK* (*Pek*), *Atf4*, or *Atf6* in either *Dome^Meso^*^+^ or *Tep4*^+^ cells does not alter lymph gland development (Figures S7A and S7B; Figures S7F–S7K).

To further explore the genetic interactions between the Ire1–Xbp1 pathway with S-NO mediated by NO, we treated larvae with the UPR inducer, dithiothreitol (DTT). Unlike 4-PBA, which reduces the numbers of progenitor cells, DTT treatment reduces the differentiation of Pxn^+^ blood cells while maintaining a constant ratio of *Dome^Meso^*^+^ progenitors (Figures 6I–6K). Additionally, the precocious differentiation phenotype caused by knockdown of *Nos^Oxy^*in *Dome^Meso+^* progenitors (*Dome^Meso^-Gal4 UAS-Nos^Oxy^ RNAi*) is restored by DTT activation of the UPR (Figures 6I–6K). Overexpression of *Ire1* in *Dome^Meso^*^+^ progenitors (*Dome^Meso^-Gal4 UAS-Ire1*) decreases Pxn^+^ blood cell differentiation and increases *Dome^Meso^*^+^ progenitor cells (Figures 6L and 6N). Furthermore, progenitor-specific expression of *Ire1* is sufficient to restore the differentiation phenotype caused by loss of *Nos^Oxy^*(*Dome^Meso^-Gal4 UAS-Ire1 UAS-Nos^Oxy^ RNAi*) (Figures 6L and 6N). In contrast, overexpression of *Nos*, which increases *Dome^Meso^*^+^ progenitor cells, does not affect the *Xbp1*-knockdown phenotype (*Dome^Meso^-Gal4 UAS-Xbp1RNAi UAS-Nos*) (Figures 6M and 6O), suggesting that the Ire1–Xbp1-induced UPR pathway functions downstream of Nos. Based on these findings, we conclude that the Ire1–Xbp1-dependent UPR pathway is crucial for core progenitor maintenance and acts downstream of S-NO.

### The S-nitrosylation/unfolded protein response pathway promotes G2-cell-cycle arrest in blood progenitors

Blood progenitors are arrested in the G2 phase after reaching the third instar^102^, but the mechanisms underlying how progenitors are maintained in the G2 cell cycle remain unclear. Given the link between the UPR and cell cycle control^103–105^, we investigated whether the NO-dependent UPR pathway alters cell cycle progression in blood progenitors. To monitor the cell cycle transition in vivo in the lymph gland, we used the Fly-FUCCI system^106^ under control of the progenitor-specific *Dome^Meso^-Gal4* driver (*Dome^Meso^-Gal4 UAS-FUCCI*). As previously reported^107, 108^, most (∼ 58%) of wild-type lymph gland progenitor cells pause in the G2 phase, while the rest remain in G1 (∼ 14%) or S phase (∼ 27%) in the third instar (Figures 7A and 7E; Figures S8A and S8B). Knockdown of *Nos^Oxy^* in the progenitors leads to a marked reduction in the proportion of cells expressing G2 (∼ 24%) (*Dome^Meso^-Gal4 UAS-FUCCI UAS-Nos^Oxy^ RNAi*) (Figures 7A and 7E; Figures S8A and S8B). This genetic background exhibits a reduced number of *Dome^Meso^*-driven *FUCCI*-positive cells, again validating that *Nos* loss induces precocious differentiation of blood progenitors. In addition, *IP3R RNAi* expression in the progenitor results in a progenitor ratio in G2 similar to that of progenitors treated with *Nos RNAi* (*Dome^Meso^-Gal4 UAS-FUCCI UAS-IP3R RNAi*) (Figures S8C–S8F). Although reducing *Nos* or calcium levels alone is sufficient to delay entry of the progenitors into the G2 phase of the cell cycle, neither an increase in calcium levels nor overexpressing *Nos* changes the cell cycle profiles of progenitor cells (*Dome^Meso^-Gal4 UAS-FUCCI UAS-IP3R* or *Dome^Meso^-Gal4 UAS-FUCCI UAS-Nos*) (Figures 7B and 7E; Figures S8A–S8F). Similarly, an increase in cytosolic calcium induced by overexpression of *IP3R* does not recover the *Nos* RNAi-mediated decrease in the ratio of G2 cell cycle (*Dome^Meso^-Gal4 UAS-FUCCI UAS-Nos RNAi UAS-IP3R*) (Figures S8C–S8F). Furthermore, overexpression of *Nos* does not rescue the reduced G2 phenotype caused by *IP3R* RNAi (*Dome^Meso^-Gal4 UAS-FUCCI UAS-IP3RRNAi UAS-Nos*) (Figures S8C–S8F). These results suggest that both NO and calcium are required for maintaining progenitor cells in G2, and neither NO nor calcium alone is sufficient to maintain progenitors in G2.

**Figure 7.**
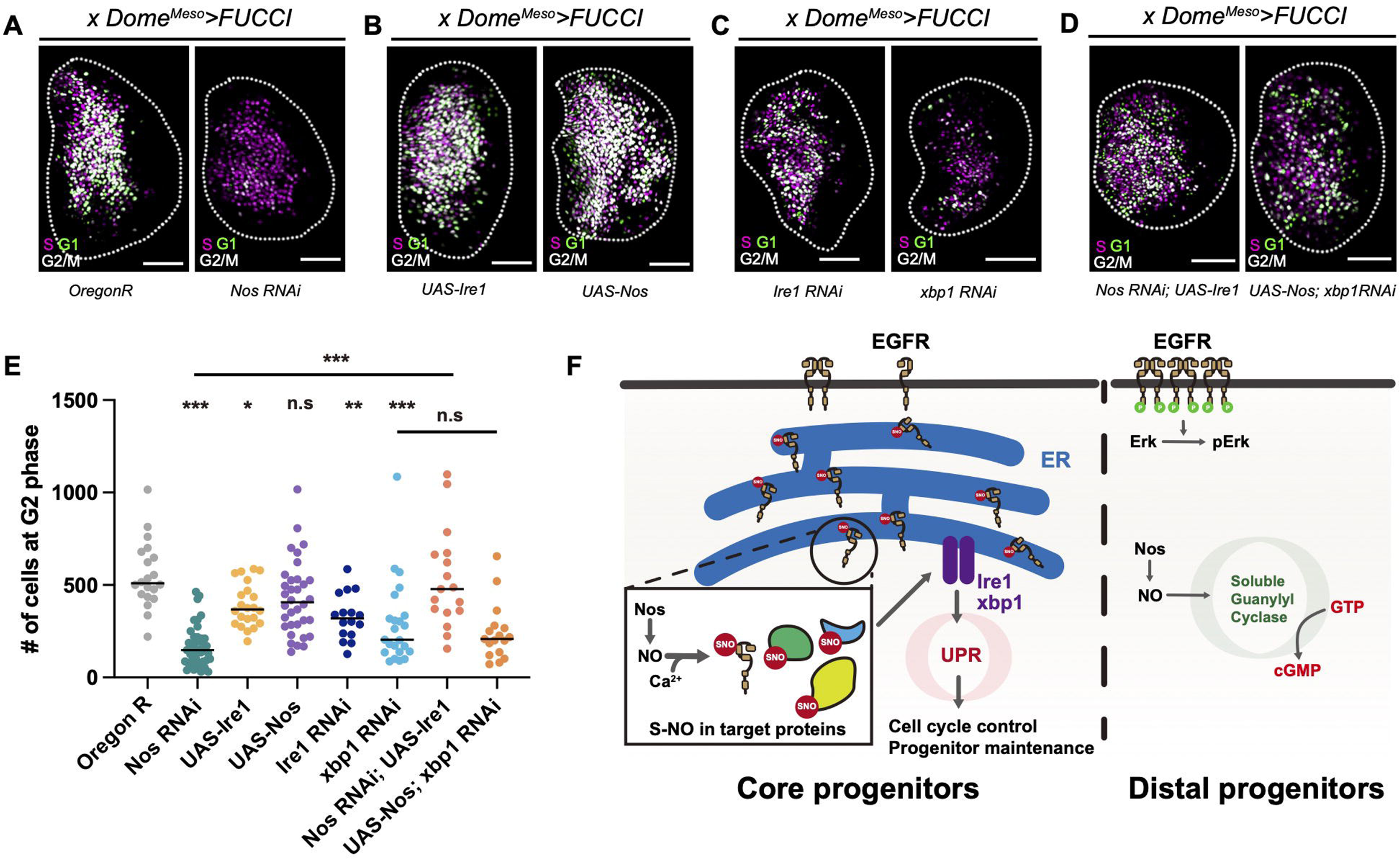
Ire1/Xbp1-dependent UPR pathway promotes the G2 cell cycle arrest. A-E. Cell cycle profiles of progenitor cells monitored by Fly-FUCCI. Progenitor-specific RNAi against *Nos* reduces the G2 phase of the cell cycle (white) in progenitor cells (*Dome^Meso^-Gal4 UAS-FUCCI UAS-NosRNAi*) **(A)**. Overexpression of *Ire1* or *Nos* does not alter G2 cell cycle arrest in the lymph gland (*Dome^Meso^-Gal4 UAS-FUCCI UAS-Ire1 or UAS-Nos*) **(B)**. Knockdown of *Ire1* or *Xbp1* in progenitor cells reduces the proportion of G2 phase (white) in the lymph gland (*Dome^Meso^-Gal4 UAS-FUCCI UAS-Ire1RNAi or UAS-Xbp1RNAi*) **(C)**. Simultaneous expression of *Ire1* with *Nos* RNAi restores the ratio of progenitor cells in the G2 cell cycle (white) (*Dome^Meso^-Gal4 UAS-FUCCI UAS-NosRNAi UAS-Ire1*), compared to the cell cycle profile caused by *Nos* RNAi (A). Overexpression of *Nos* with *Xbp1* RNAi does not alter the G2 cell cycle arrest (*Dome^Meso^-Gal4 UAS-FUCCI UAS-Xbp1RNAi UAS-Nos*) **(D)**. Quantification of the number of cells in the G2 phase shown in (A-D) **(E)**. *Oregon R* (n=46), *Nos RNAi* (n=40), *UAS-Ire1* (n=22), *UAS-Nos* (n=17), *Ire1 RNAi* (n=22), *Xbp1 RNAi* (n=23), *UAS-Nos; UAS-Xbp1RNAi* (n=17)*, UAS-Ire1; UAS-NosRNAi* (n=17). **F.** Schematic illustration of NO-mediated control of blood progenitor cells in the lymph gland. In core progenitors (Col^+^, *Tep4*^+^, *Dome^Meso+^*) (left), S-NO blocks EGFR trafficking, induces the *Ire1*/*Xbp1*-dependent UPR pathway, and maintains the progenitor cell fate. In distal progenitors (Col^-^, *Tep4*^-^, *Dome^Meso+^, Nplp2^+^*) (right), EGFR is activated and induces dpErk. Free-radical NO induces soluble guanylyl cyclase to generate cGMP. White scale bar, 30 μm; yellow scale bar, 3 μm. White dotted lines demarcate the lymph gland. n.s.: not significant (p>0.01); \**p*<0.01; \*\**p*<0.001, \*\*\**p*<0.0001. Bars in graphs: the median.

Consistent with the above findings, we observed a similar reduction in the G2 cell cycle by knocking down either *Ire1* or *Xbp1* (*Dome^Meso^-Gal4 UAS-FUCCI UAS-Ire1 RNAi* or *Dome^Meso^-Gal4 UAS-FUCCI UAS-Xbp1 RNAi*) (Figures 7C and 7E; Figures S8A and S8B). Similar to *Nos*, overexpression of *Ire1* alone in *Dome^Meso^*^+^ progenitors does not alter the ratio of G2 cell cycle in the blood progenitors (*Dome^Meso^-Gal4 UAS-Ire1*) (Figures 7B and 7E; Figures S8A and S8B). Notably, co-expression of *Ire1* with *Nos^Oxy^* RNAi is sufficient to restore the ratio of cells in the G2 phase compared with *Nos^Oxy^* RNAi alone (*Dome^Meso^-Gal4 UAS-FUCCI UAS-Ire1 UAS-Nos^Oxy^ RNAi*) (Figures 7D and 7E; Figures S8A and S8B). However, overexpression of *Nos* with *Xbp1* RNAi in *Dome^Meso^*^+^ progenitors does not rescue the reduction in cells in the G2 phase caused by *Xbp1* RNAi (*Dome^Meso^-Gal4 UAS-FUCCI UAS-Xbp1 RNAi UAS-Nos*) (Figures 7D and 7E; Figures S8A and S8B). Taken together, these findings suggest that the Ire1–Xbp1-dependent UPR pathway is required for G2 cell cycle in the blood progenitors downstream of S-NO to maintain blood progenitor cell fates.

## Discussion

In this study, we investigated the role of S-NO, a protein modification facilitated by NO, in the maintenance of blood progenitor cell fates in the lymph gland. Our findings showed that NO expression in core progenitors triggers S-NO of proteins in blood progenitors in combination with cytosolic calcium, enhancing the Ire1–Xbp1-mediated UPR necessary for maintaining blood progenitors. We discovered EGFR as a potent target of S-NO and demonstrated that its translocation to the cell membrane is inhibited by S-NO during progenitor maintenance. These results provide new insights into how blood progenitors confine the spatio-temporal expression of critical proteins through NO-mediated post-translational modifications and into the role of a developmentally controlled UPR in blood stem/progenitor cell maintenance.

The multifaceted functions and regulatory pathways associated with EGFR in development and tumorigenesis have been studied extensively; however, the mechanisms that determine the amount of EGFR at the cell membrane in the absence of ligands at a given time or space remain largely unknown. In the lymph gland, EGFR is a well-known upstream receptor for several signaling cascades, including the Ras/MAPK pathway, which stimulates the proliferation and differentiation of blood progenitors^109^. Therefore, lymph gland progenitors keep EGFR signaling low to maintain blood progenitor cell fates prior to the onset of differentiation. S-NO of EGFR may be a key mechanism that induces EGFR dormancy in the ER; EGFR dormancy can be relieved immediately during stress responses, such as immune activation, to allow proper protein translocation and cell differentiation. Interestingly, the two cell-autonomous factors that determine the level of S-NO, cytosolic calcium and NO, integrate environmental information into the core progenitors. Cytosolic calcium levels in blood progenitors are regulated by various membrane receptor proteins, including GABA_B_ receptors^46^. Similarly, the expression of NO is modulated by environmental changes, such as ambient oxygen levels^80^. Thus, the levels of calcium and NO together may provide environmental information to progenitor cells to determine which cells differentiate and when. The expression of EGFR^C311S^ inhibits EGFR localization to the cell membrane in the salivary gland and lymph gland progenitors; therefore, the control of membrane translocation via S-nitrosylated EGFR is likely conserved in other cell types. In addition to EGFR, NO targets additional proteins involved in blood progenitor maintenance, such as cell cycle regulators. Accordingly, validation of additional S-NO protein targets in stem/progenitor maintenance will be critical to understanding the significance of S-NO and protein localization control in cell fate determination.

Nos is expressed in the blood progenitor cells of the lymph gland, but it lacks the reductase domain found in typical Nos enzymes. Despite this deficit, Nos can still generate NO and initiate downstream signaling cascades. Interestingly, lymph gland progenitors express reactive oxygen species (ROS) as a developmental signal to maintain progenitor cell fate^81, 110^. Nos^Oxy^ in blood progenitor cells may incorporate ROS to synthesize NO in the absence of the reductase domain. This interplay between ROS and NO may help maintain appropriate levels of ROS, generate NO, and protect progenitor cells from excess free radical expression. We showed that NO generated in the core progenitors is primarily utilized for S-NO, but a subset of cells in the IZ expresses the free radical form of NO to activate cGMP-mediated signaling, possibly via soluble sGC, a potent NO sensor. Future studies will investigate how the sGC-cGMP signaling cascade controls the development of blood progenitors in the IZ or maturation of plasmatocytes or crystal cells in the CZ.

The rate of protein translation influences stem cell differentiation in vertebrates^111^. Furthermore, previous studies have shown that the UPR, a stress response mechanism triggered by protein misfolding, is implicated in regulating clonal proliferation of hematopoietic stem cells in response to various stress signals^112^. Protein synthesis represents a potential convergence point for stress responses and developmental signals, and the UPR can override several upstream regulators that modify protein translation or trafficking. The UPR pathway is well conserved between *Drosophila* and humans and has been linked to tissue homeostasis and lifespan regulation in *Drosophila*^26^. Recent studies in *Drosophila* have also implicated the UPR in the proliferation and regenerative capacity of intestinal stem cells^23,113,114^. Because intestinal stem cells are significantly influenced by ROS and NO, it would be interesting to investigate whether the ROS-NO induced UPR is a core regulatory mechanism in stem/progenitor cells. Unlike *Nos RNAi*, inhibition of the UPR significantly reduces the expression of *Dome^Meso+^* progenitors and causes progenitor cells to lose both progenitor and differentiating blood cell markers (Figures 7A–7F). This phenotype suggests functional divergence of the UPR pathway in the stem/progenitor control independent of S-NO^115^. Taken together, these findings highlight that the UPR pathway can be endogenously activated by developmental programs, such as S-NO, in stem/progenitor cells to promote determination of their fates in the lymph gland.

## Acknowledgments

The authors thank the members of the Shim lab for helpful discussions. The authors acknowledge the Bloomington, VDRC, DGRC, NIG, and KDRC *Drosophila* stock centers and the DSHB hybridoma bank. The authors thank the following individuals for stocks and reagents: Drs. I. Ando, U. Banerjee, M. Crozatier, P. O’Farrell, M.J. Kang, S. Lee, O. Schuldiner, and S. Sinenko. This study was supported by grants from the National Research Foundation (NRF) of Korea (NRF-2022R1I1A1A01069677) to B.C. and (NRF-2019R1A2C2006848 and RS-2023-00218602) to J.S.

## Author contributions

Conceptualization: J.S. and B.C.; Methodology: B.C., M.S., E.C., and S.S.; Investigation and analysis: B.C., M.S., E.C., and S.S.; Data curation: B.C., M.S., and J.S.; Writing: B.C., M.S., U.B., and J.S.; Funding acquisition and supervision: J.S.

## Declaration of competing interests

The authors declare no competing interests.

## STAR Methods

### Drosophila stocks and genetics

The following *Drosophila* stocks were used in this study: *Dome^Meso^-Gal4* (U. Banerjee)*, Hml*^Δ^*-Gal4* (S. Sinenko)*, Tep4-Gal4* (U. Banerjee)*, Lz-Gal4* (U. Banerjee), *Nos^Oxy^ RNAi* (BL28792, NIG6713R)*, Nos^Red^ RNAi* (V27722)*, UAS-Nos* (BL56830)*, Ire1 RNAi* (V39561), *Xbp1 RNAi* (BL36755), *Pek RNAi* (BL35162), *crc RNAi* (BL80388), *Atf6 RNAi* (BL26211), *Atf6-GFP.FPTB* (BL83413), *hs-flp, Actin-FRT-Stop-FRT-Gal4,UAS-EGFP* (U. Banerjee)*, EGFR-sfGFP* (BL92329)*, Nplp2-Gal4* (KDRC10023)*, UAS-GCaMP3* (BL32237)*, UAS-RcaMP* (BL51928), *HHLT-Gal4* (U. Banerjee)*, Ubi-Gal4* (U. Banerjee), *IP3R RNAi* (BL25937)*, UAS-IP3R* (U. Banerjee), *UAS-Ire1* (KDRC 10875)*, UAS-LacZ* (U. Banerjee), *Nos^1^* (BL56822), *Nos*^Δ*1*5^ (P. O’Farrell)^83^*, Nos*^Δ*all*^ (O. Schuldiner)^82^*, UAS-FUCCI* (BL55122)*, UAS-Xbp1::GFP* (BL39720)*, UAS-mCherry* (BL52268)*, UAS-KDEL-GFP* (BL9898)*, UAS-KDEL-RFP* (BL30909), and *Oregon R* (BL5).

Generation of *UAS-FLAG::EGFR^WT^* or *UAS-FLAG::EGFR^C311S^*flies: A vector with an N-terminal-FLAG-tagged *EGFR* sequence was a gift from S. Lee^116^. For subcloning, we used the *FLAG::EGFR* vector as a template with using the following primers.

Forward: 5’-GGGCTCGAGATGCTGCTGCGACGGCGCAA-3’

Reverse: 5’-GGGTCTAGACTACACCCTCGTCT CGTGTTGCGG-3’

To create the EGFR^C311S^ mutation, we used a mutagenesis kit (EZ004S; Enzynomics, Inc., Daejeon, South Korea) and followed the manufacturer’s instructions. The following primers were used for the mutagenesis.

Forward: 5’-GACTGGTCCCACGCAAAAG-3’

Reverse: 5’-GATCCTCCGGCGCAGAAGA-3’

DNA was subcloned into the pUASt-AttB vector (DGRC 1419). Transgenic flies were generated by BestGene Inc. (Chino Hills, CA, USA).

All fly stocks were maintained at 25°C, except for the heat shock clone experiment. To generate heat shock clones, crossed flies were kept at 18°C until the second instar. At the second instar, larvae-containing vials were heated in a 37°C water bath for 1 h. After heat shock, vials were placed at 18°C again and transferred to 25°C after a day until larvae reached the desired developmental stage.

### Immunohistochemistry

Lymph glands were dissected and stained as previously described^52^. Briefly, lymph glands were fixed in a 3.7% formaldehyde solution and washed three times using 0.4% PBS TritonX-100. After three washes, samples were blocked using 1% BSA for 30 min at RT (25°C). Samples were incubated with the desired primary antibodies overnight at 4°C and then washed three times using 0.4% PBS TritonX-100. Samples were incubated with secondary antibodies (Cat. #: 115-165-166, 711-165-152, 115-095-062, 711-095-152, 715-605-151, and 711-605-152; Jackson ImmunoResearch Labs, West Grove, PA, USA) diluted 1:250 for 3 h. Samples were then washed three times. Finally, samples were maintained in VECTASHIELD (Vector Laboratories, Inc., Newark, CA, USA) until they were mounted on glass slides.

The following primary antibodies were used: □-Hnt (DSHB 1G9, 1:10), □-Pxn (1:1000)^117^, □-collier (Gift from M. Crozatier, 1:400), □-NimC1 (Gift from I. Ando, 1:100), □-Nos (Gift from O. Schuldiner, 1:100), □-SNO (Abcam 94930, 1:100), □-FLAG (Sigma F1804, 1:1000), □-GFP-Rb (Abcam 290, 1:1000), □-GFP-Mouse (Sigma G6539, 1:1000), □-crc (Gift from M.J. Kang)^25^, □-HA-Rabbit (Cell signaling 3724S, 1:100), and □-dpErk (Cell signaling, 4370S, 1:100). Images were obtained with a Nikon C2 Si-plus confocal microscope.

### Western blotting

To extract proteins from the lymph gland, a minimum of 100 lymph glands were prepared in Schneider’s medium for each experiment. After preparation, samples were centrifuged at 4°C, 3000 rpm for 5 min, and lymph gland pellets were lysed with 50 µl RIPA buffer (MB-030-0050, Rockland) containing a protease inhibitor cocktail (P9599, Sigma). Lysed samples were centrifuged at 4°C, 13,000 rpm for 15 min, and supernatants were used for the experiment. At least 20 µg of proteins were loaded onto the gel. The following antibodies were used: □-Nos (Gift from O. Schuldiner, 1:1000), □-GFP-Rb (Abcam 290, 1:1000), □-GFP-Mouse (Sigma G6539, 1:1000), □-Erk (Cell signaling 9102S, 1:1000), □-dpErk (Cell signaling 4370S, 1:1000), □-Tub (DSHB 12G10, 1:1000), □-Mouse-HRP (Cell Signaling 7076S, 1:1000), □-Rabbit-HRP (Cell Signaling 7074S, 1:1000), □-Guinea Pig-HRP (Bethyl A60-110P, 1:1000). Images were visualized using an HRP substrate (Sigma, WBLUF0100) and captured by VILBER Fusion SL or by Medical X-ray FILM (AGFA, CP-BU 810).

### iodoTMT-labeling of S-nitrosylated proteins

For iodoTMT-labeling of the S-nitrosylated proteins in the lymph gland, we used the Pierce S-Nitrosylation Western Blot Kit (90105, Thermo Fisher Scientific) and followed the manufacturer’s procedure. Briefly, proteins extracted from 1000 lymph glands were treated with MMTS that block un-nitrosylated cysteines. After blocking, proteins were precipitated using 6 volumes of cold acetone (1720°C) and kept at 1720°C for 1 h. Precipitated proteins were centrifuged at 10,000 g for 10 min at 4°C. Supernatants were removed, and protein pellets were dried for 10 min at RT and re-suspended in 50–100 ul HENS buffer. For a negative control, sodium ascorbate-treated and untreated samples were prepared and added to the iodoTMT-labeling reagent. Samples were maintained for more than 2 h at RT and then used in the desired experiments.

### Chemical feeding

For inducing ER stress in the lymph gland, larvae were grown in 5 mM DTT (10197777001, Sigma) with normal cornmeal food for 24 h before dissection. For 4-PBA (P21005, Sigma) experiments, larvae were grown in 4 mM of 4-PBA-containing normal cornmeal food for 48 h before dissection.

### Quantification of samples

For all experiments, more than three independent biological replicates were conducted. Stained or fluorescently labeled cells were quantified and analyzed by Image J (NIH, USA) or IMARIS 8.3 software (Bitplane, Belfast, UK). Individual primary lobes were counted. In vivo data were analyzed by the Mann–Whitney test after determining normality using SPSS (version 24).

### DAF-FM diacetate (free radical NO) and DHE (ROS) staining

Larvae were dissected at 96 h AEL in Schneider’s medium and kept on an ice-cold glass well plate during dissection. After dissection, the medium was replaced with 1:1000 diluted DAF-FM diacetate (D23844, Invitrogen) or DHE (D11347, Invitrogen) medium. For the DAF-FM diacetate and DHE incorporation, samples were shaken for 10 min at RT. Samples were washed three times in RT Schneider’s medium for 3 min. After washing, samples were fixed in 3.7% formaldehyde for 3 min. To remove the fixative solution, samples were washed with 1X PBS for 3 min. Samples were then mounted with VECTASHIELD (Vector Laboratories) without DAPI and imaged using a Nikon C2 Si-plus confocal microscope.

### Immunoprecipitation and LC–MS

For immunoprecipitation, 1000 lymph glands were prepared in Schneider’s medium and centrifuged at 4°C, 3000 rpm for 5 min and repeated this 10 times. Lymph gland pellets were kept at 1780°C until the desired number of lymph glands were collected. For each experiment, 1000 lymph glands were needed. For the LC–MS experiment, 10,000 lymph glands were prepared. When samples were ready, Protein A agarose beads (20333, Thermo Fisher Scientific) and the desired antibody (L-GFP; Sigma G6539) were prepared in 1 ml of pH 7.5, 50 mM Tris-HCl buffer. Agarose beads with antibodies were incubated at 4°C overnight with rotation. During the antibody-to-agarose conjugation step, 1 ml lysis buffer (pH 7.5, 250 mM Tris-HCl buffer, 250 mM NaCl, 1.5 M sucrose, 1% TritonX-100, 1X protease inhibitor cocktail (P9599, Sigma), 0.2 mM PMSF) was used to lyse the lymph gland samples. Lysed samples were centrifuged at 4°C, 12,000 g for 15 min. One milliliter of protein lysate was transferred to a tube containing antibody-agarose conjugate and incubated at 4°C overnight with rotation. The day after incubation, samples were placed onto micro spin columns (7326204, BioRad) and washed three times with wash buffer (pH 7.5, 250 mM Tris-HCl buffer, 250 mM NaCl, 1.5 M sucrose, 0.2% TritonX-100, 1X protease inhibitor cocktail (P9599, Sigma), 0.2 mM PMSF) using gravity flow. Finally, 40 µl of sample and 2X SDS sample buffer were boiled and collected by centrifugation. Protein samples were loaded onto a 7.5% of SDS–PAGE gel and stained with Coomassie Brilliant Blue solution (JB1000, Taeshin Bio). LC–MS was performed at the Yonsei Proteome Research Center (Seoul, South Korea).

### Quantitative real-time PCR analysis

At least 100 larval lymph glands were dissected to extract RNA. cDNA was synthesized using the ReverTra ACE qPCR RT Master Mix (TOFSQ-201, TOYOBO). qRT-PCR was performed by using the PowerUp SYBR Green Master Mix (A25741, Thermo Fisher) and a StepOnePlus Real-Time PCR System (Applied Biosystems). The relative quantity of the target gene was calculated using the comparative ΔΔCt method. The following primers were used.

BiP Forward: 5’-TGTCACCGATCTGGTTCTTCAGGC-3’

Bip Reverse: 5’-GTCCCATGACCAAGGACAACCATC-3’

Pdi Forward: 5’-AGATCTCGTCCTTCCCCACA-3’

Pdi Reverse: 5’-TCGAGAGTCCTGTCCAGGTT-3’

Ire1 Forward: 5’-ATGGTAAGGAGGGCGAGCAG-3’

Ire1 Reverse: 5’-ATGACCGTGTACTGAGTC-3’

Xbp1 Forward: 5’-AGGCCATCAACGAGTCACTG-3’

Xbp1 Reverse: 5’-CTTTCCAGAGTGAGGCCAGG-3’

### SABER in situ hybridization

All procedures were conducted as described previously^86^, except for a few modifications. Lymph glands were dissected in Schneider’s medium within 30 min and fixed in 4% formaldehyde for 30 min. After fixation, samples were washed with 0.3% PBST (Tween-20) three times for 5 min each. Samples were washed at least twice for 60 min with wash hybridization buffer (wHyb) and treated with two different primary probes for 48 h at 43°C. After probe hybridization, samples were washed with wHyb or 2X SSCT buffer and treated with imaging probes. To amplify the signal, the HRP imaging probe was used instead of the Cy3 imaging probe. To boost the HRP signal, the SuperBoost Kit (B40933, Thermo Fisher) was used. Samples were maintained in VECTASHIELD until mounted on glass slides. Probes used in this study are listed below.

*Nos oxidase*: 5’-CTGTCGACATGTACCAACCACTGAATGTGTTTCATCATCAT-3’, 5’-CGGGCGGGTAATCAAAGTAGTCCGGATCTTTCATCATCAT-3’

*Nos reductase*: 5’-GCATCGTTGTCTTGGCTCCCTCACTCAAACTTTCATCATCAT-3’, 5’-CTTGGGCAATTTGGCCGTGGGATCAAGTTTCATCATCAT-3’

### Data availability

All data generated during and/or analyzed during the current study are included in this published article and its supplementary information files.

---

**Sup Figure 1.**
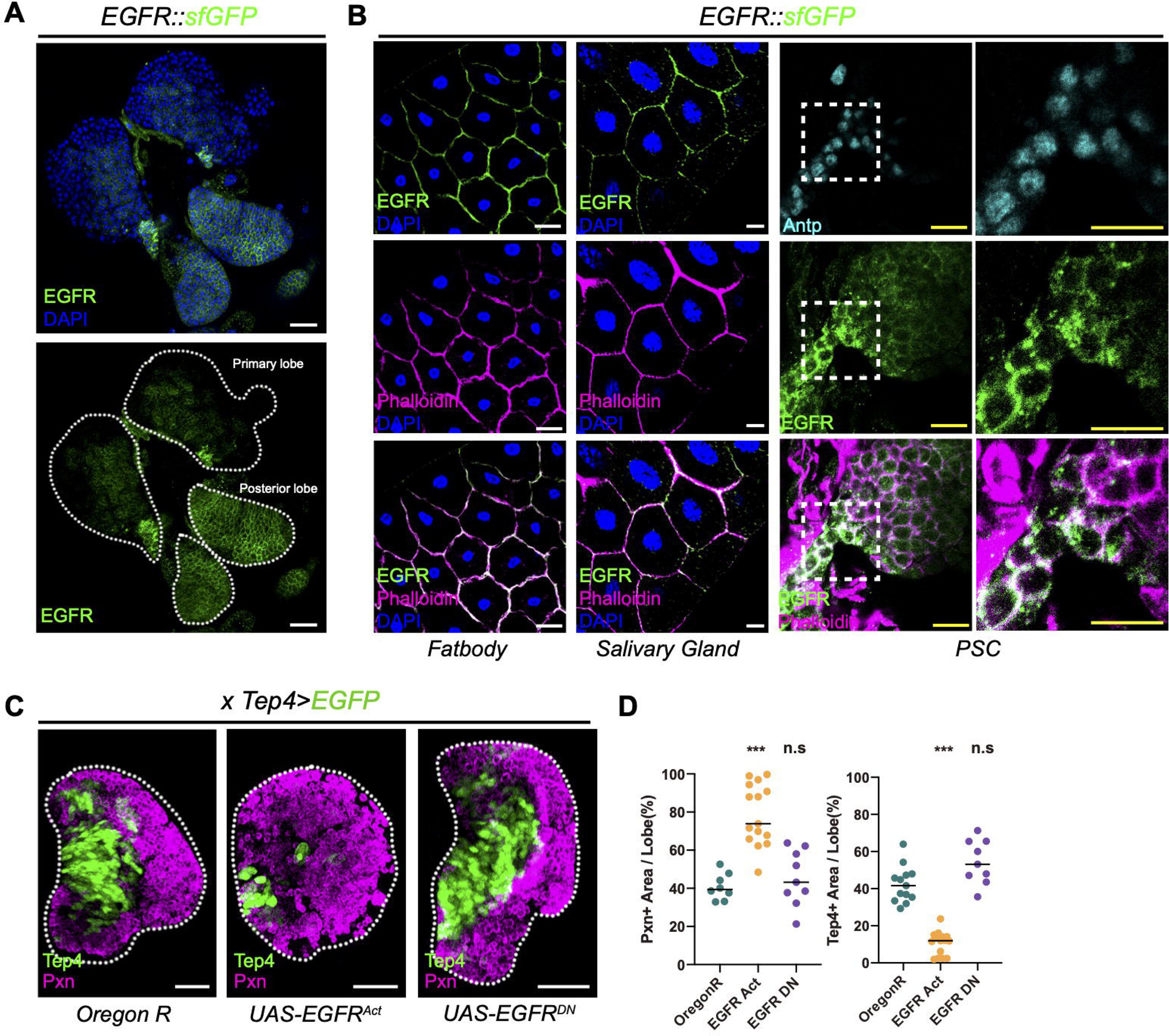

**Sup Figure 2.**
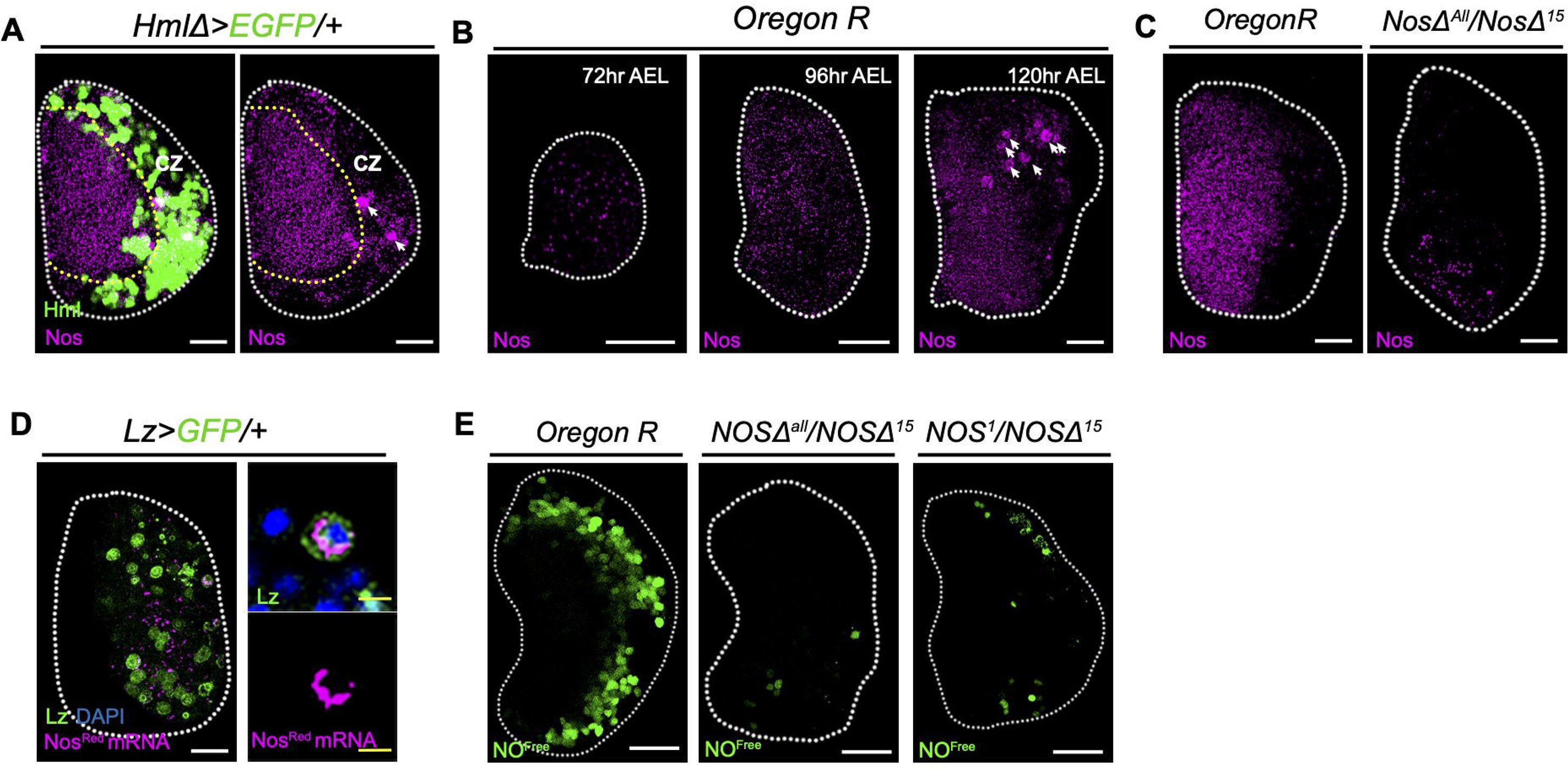

**Sup Figure 3.**
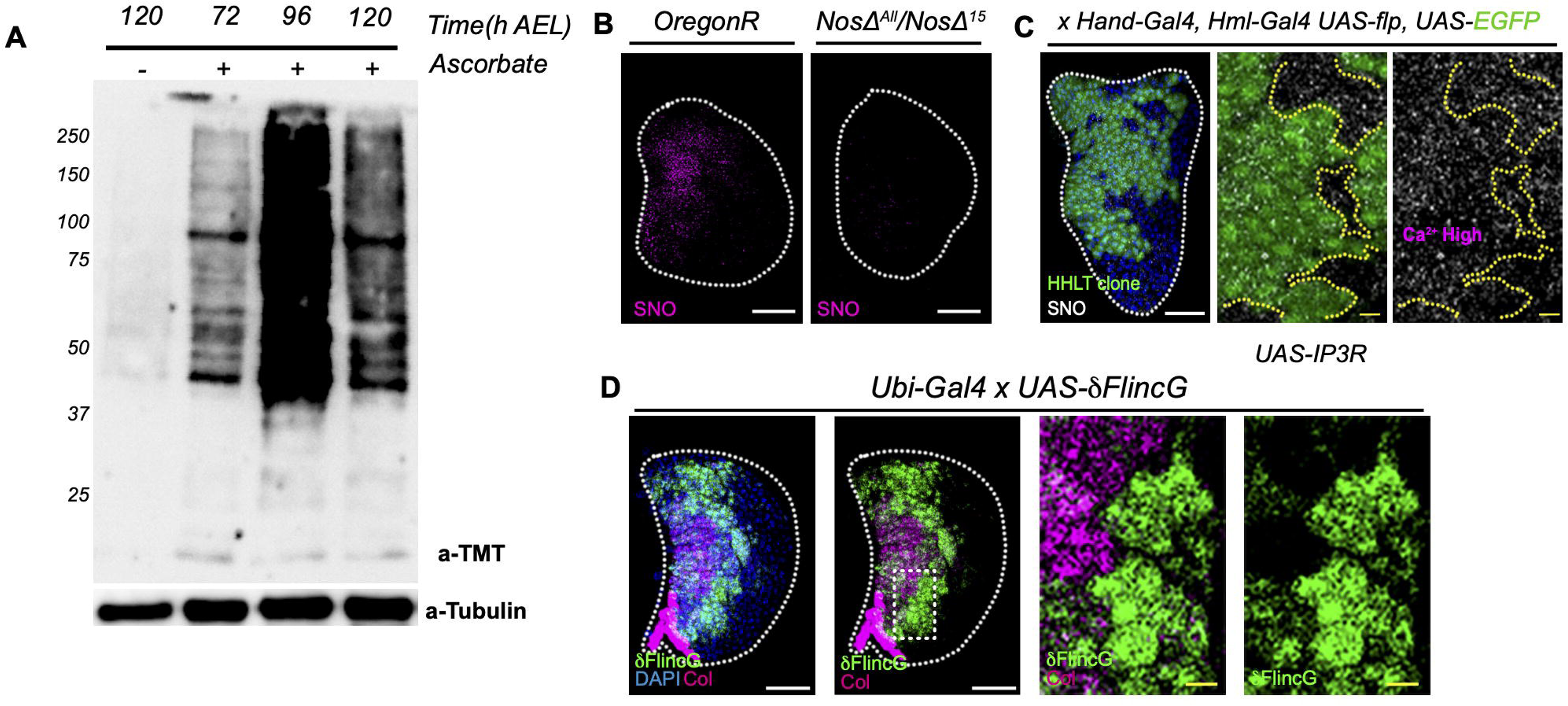

**Sup Figure 4.**
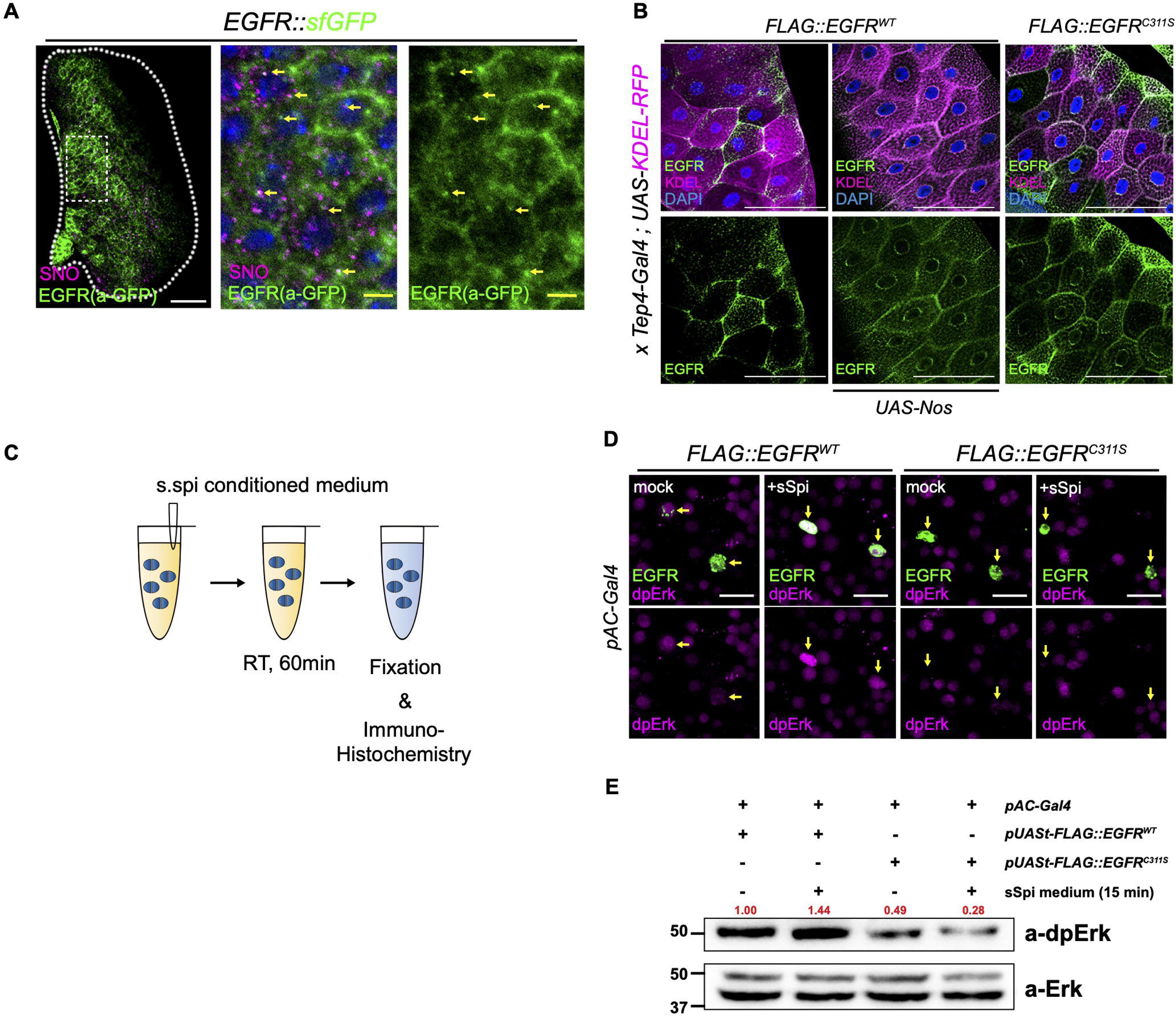

**Sup Figure 5.**
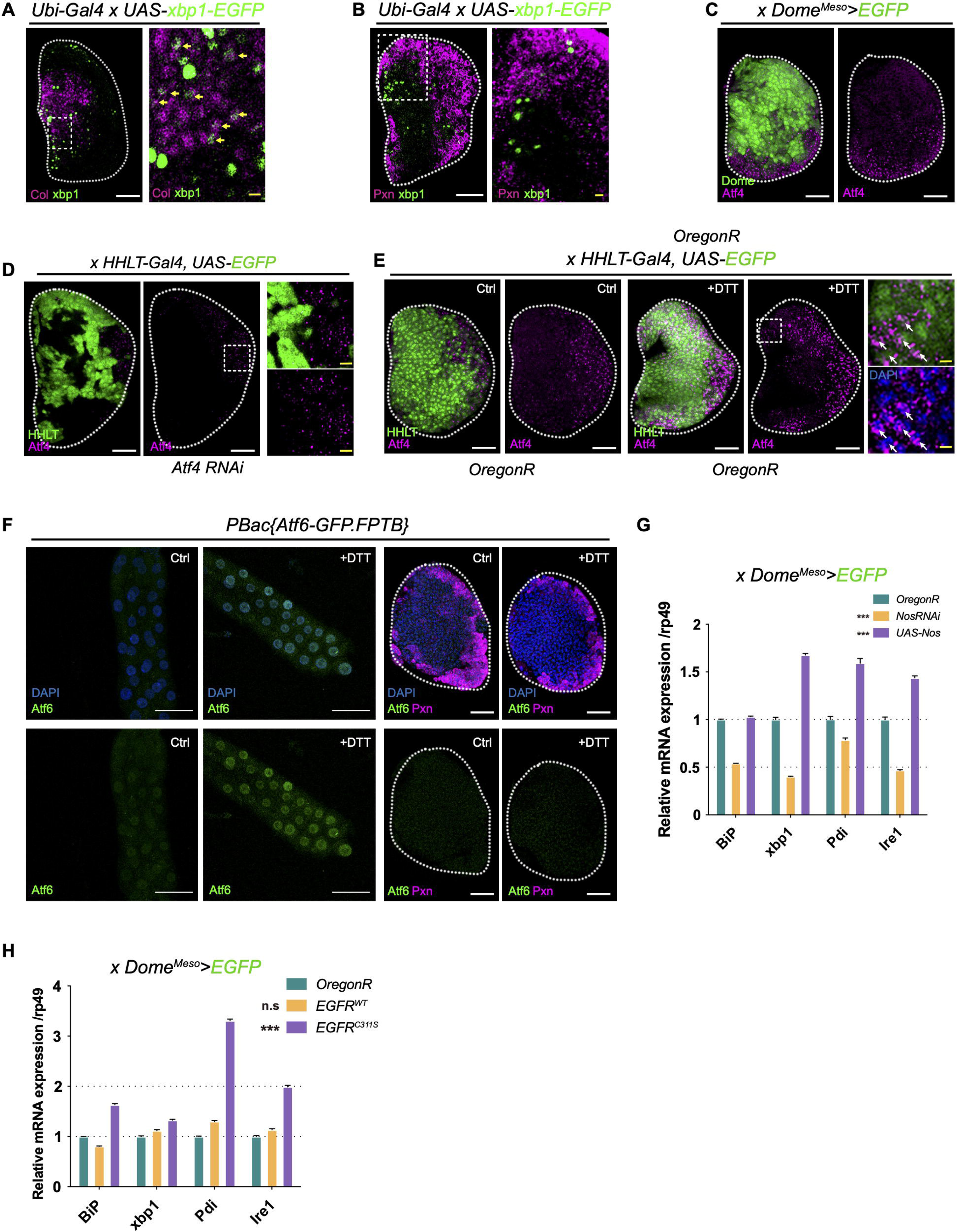

**Sup Figure 6.**
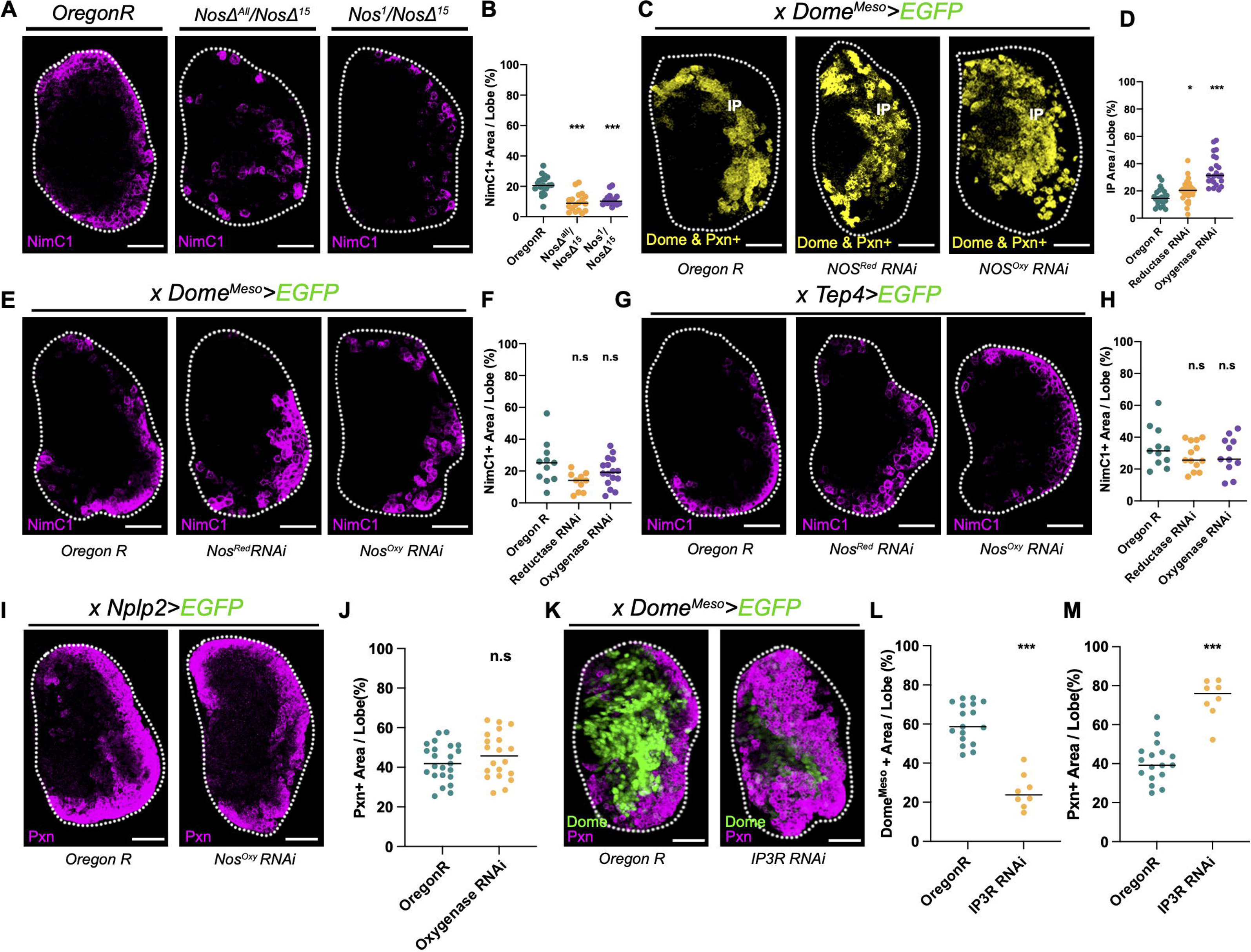

**Sup Figure 7.**
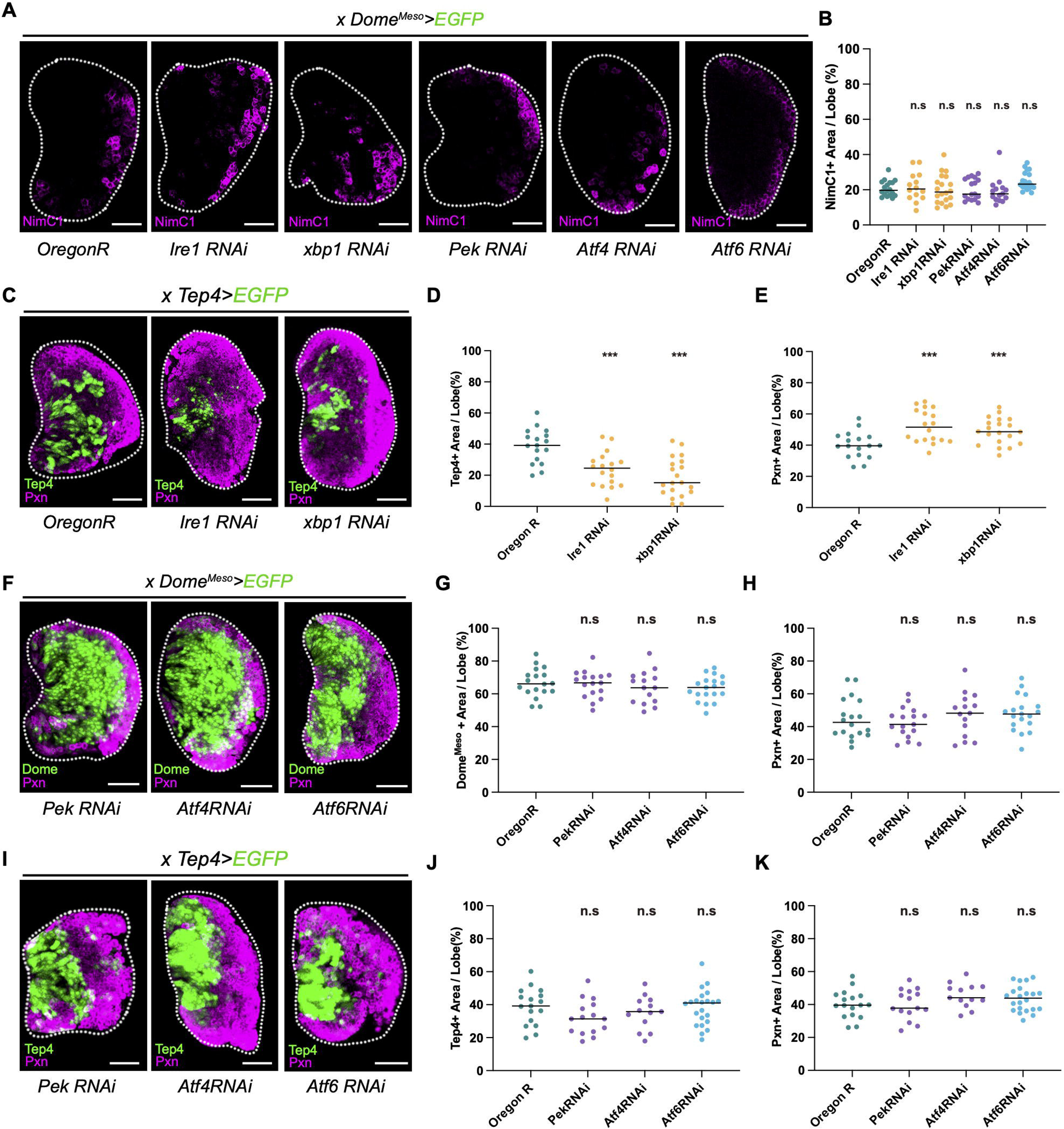

**Sup Figure 8.**
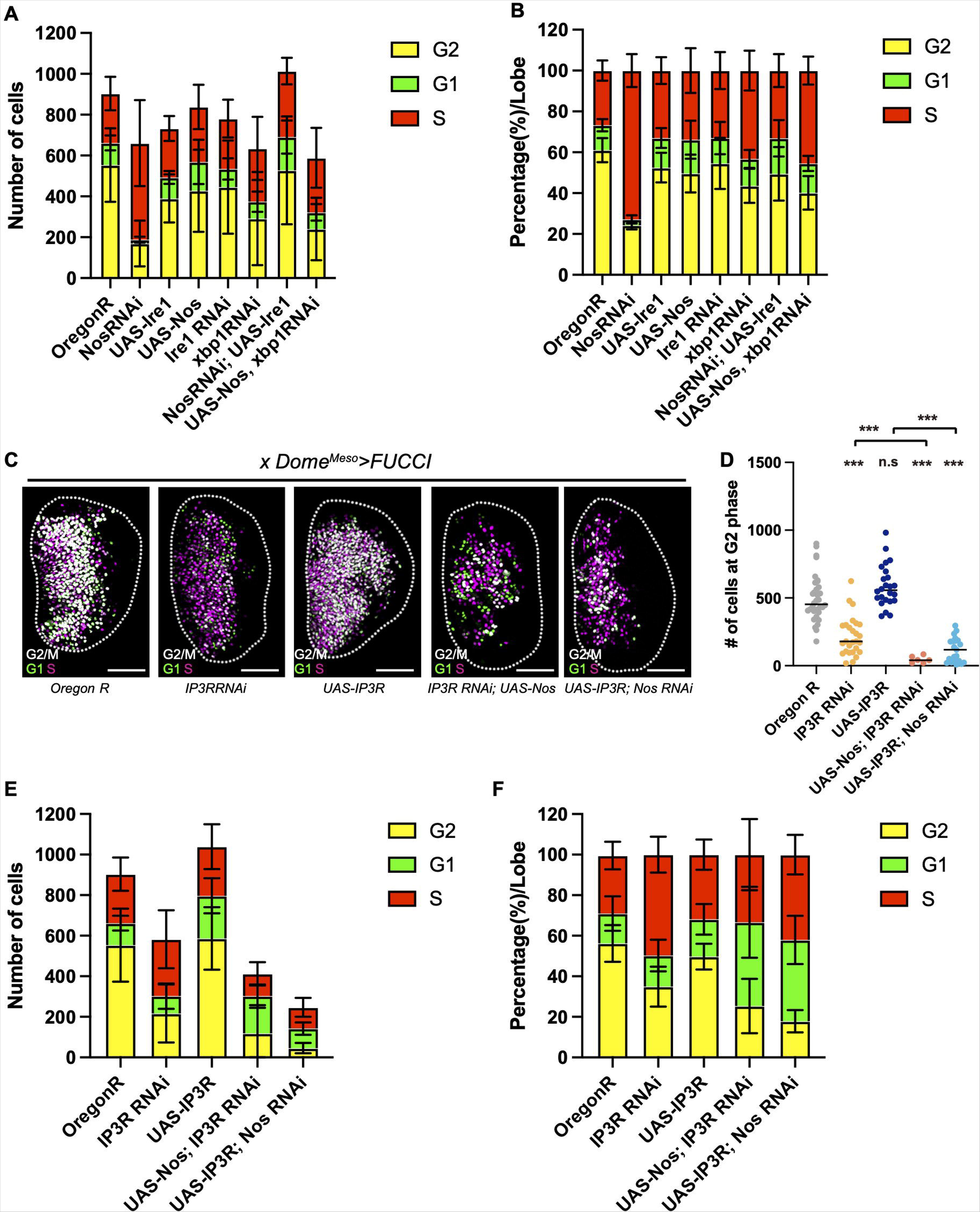

## Notes

### Competing Interest Statement

The authors have declared no competing interest.

