## Supplemental Figure Legend for "S-nitrosylation-mediated unfolded protein response maintains hematopoietic progenitors"

**Supplementary Figure Legends**

**Supplementary Figure 1. Functional attenuation of EGFR in the lymph gland**

**A.** Expression of EGFR (green) in the lymph gland primary and posterior lobes (*EGFR-sfGFP*).

**B.** EGFR (green) co-localizes with the membrane-specific marker, phalloidin (magenta), in the fat body (left), salivary gland (middle), or Antp^+^ PSC (cyan) (right). Magnified images of the PSC (inset) are shown on the right.

**C-D.** Expression of constitutive active (*EGFR^Act^*) (middle) or dominant-negative (*EGFR^DN^*) (right) form of EGFR in *Tep4^+^* progenitors (*Tep4-Gal4 UAS-EGFP UAS-EGFR^Act^* or *Tep4-Gal4 UAS-EGFP UAS-EGFR^DN^*). Compared to wild-type controls (left), *EGFR^Act^* reduces progenitor cells (green) and induces Pxn^+^ differentiating blood cells (magenta) (middle). *EGFR^DN^* does not alter lymph gland development (right). Quantification of the proportion of *Tep4*^+^ progenitors (green) or Pxn^+^ differentiating blood cells (magenta) in the lymph gland is shown in **(C-D)**.

White scale bar, 30 μm. White dotted lines demarcate the lymph gland. n.s.: not significant (p>0.01); ****p*<0.0001. Bars in graphs: the median.

**Supplementary Figure 2. Expression of Nos protein, *Nos* mRNA and free-radical NO in the lymph gland**

**A.** Nos protein expression in the lymph gland. Nos (magenta) and *Hml-*expressing differentiating blood cells (green) are mutually exclusive (*Hml^Δ^-Gal4 UAS-EGFP*). Yellow dotted line demarcates lymph gland progenitors. CZ indicates the cortical zone. White arrows indicate the crystal cell.

**B.** Nos expression during larval development. Nos (magenta) is observed at 72, 96, or 120 h AEL. White arrow indicates the crystal cell.

**C.** Specificity of the anti-Nos antibody in the lymph gland. Hetero-allelic combination of *Nos* mutants (*NosΔ^All^/NosΔ^15^*) eliminates anti-Nos expression (magenta) (right).

**D.** Visualization of *Nos* mRNA by SABER-FISH method. Reductase domain of *Nos* mRNA (magenta) is expressed in *Lz^+^* crystal cells (green) (*Lz-Gal4 UAS-GFP*). A magnified image of the crystal cell is shown on the right.

**E.** Visualization of the free radical form of NO in *Nos* mutants. Hetero-allelic combinations of *Nos* mutants (*NosΔ^All^/NosΔ^15^* or *Nos^1^/NosΔ^15^*) do not express free-radical NO as monitored by DAF-FM diacetate (NO^Free^; green).

**Supplementary Figure 3. Expression of S-NO and sGC-cGMP signaling pathway in blood progenitors**

**A.** Western blotting of TMT-labeled lymph gland proteins at three developmental time points (72, 96, and 120 h AEL). The first lane shows proteins from 120 h AEL lymph gland without ascorbate treatment during TMT labeling. Numbers on the left indicate the size of the marker proteins (kDa). Proteins were stained with anti-TMT (top) or anti-tubulin (bottom).

**B.** Visualization of S-nitrosylated proteins by anti-SNO antibody (SNO; magenta) in wild-type controls (left) or *Nos* hetero-allelic combination mutants (*NosΔ^All^/NosΔ^15^*) (right).

**C.** Expression of S-nitrosylated proteins in lymph gland clones expressing *IP3R* (green) (*Hand-Gal4, Hml-Gal4 UAS-flp, UAS-EGFP UAS-IP3RRNAi)*. Compared to the wild type (GFP-negative), clones expressing *IP3R* (green; yellow dotted line) show similar SNO (white) expression.

**D.** Visualization of cGMP signaling by a cGMP indicator, δFlincG (*Ubi-Gal4 UAS- δFlincG*). Strong δFlincG (green) and col^+^ progenitors (magenta) are mutually exclusive in the lymph gland. Magnified images are shown on the right.

**Supplementary Figure 4. S-nitrosylated EGFR inhibits its downstream signaling cascade**

**A.** Verification of S-nitrosylated EGFR by anti-SNO immunostaining. Cytoplasmic EGFR (green) co-localizes with S-NO (magenta). Magnified images are shown on the right. Yellow arrows indicate S-nitrosylated EGFR. DAPI, blue.

**B.** Localization of EGFR protein in the salivary gland. Wild-type EGFR protein is mainly localized to the membrane (*Tep4-Gal4 UAS-KDEL-RFP UAS-FLAG::EGFR^WT^*) (left). Overexpression of *Nos* with wild-type EGFR (*Tep4-Gal4 UAS-KDEL-RFP UAS-FLAG::EGFR^WT^ UAS-Nos*) (middle) or expression of EGFR^C311S^ (*Tep4-Gal4 UAS-KDEL-RFP UAS-FLAG::EGFR^C311S^*) (right) enhances EGFR expression in the ER (KDEL, magenta) and cytoplasm. EGFR protein is stained with anti-FLAG antibody (green). DAPI, blue.

**C.** Schematic representation of an ex vivo lymph gland culture with s.Spi.

**D-E.** Expression of dpErk in S2R^+^ cells. Wild-type EGFR expression in S2R^+^ cells (green; *pAC-Gal4 UAS-FLAG::EGFR^WT^*) induces dpErk (magenta) upon s.Spi treatment (left). Expression of EGFR^C311S^ (green; *pAC-Gal4 UAS-FLAG::EGFR^C311S^*) does not activate dpErk in the presence of s.Spi (right). Yellow arrows indicate S2R^+^ cells that express EGFR. Mock indicates control S2R^+^ cells without s.Spi treatment **(D)**. Quantification of dpErk activity by Western blotting **(I)**. The relative intensity of dpErk is shown in red. 37 or 50 represents the size of the protein marker (kDa).

White scale bar, 30 μm; yellow scale bar, 3 μm. White dotted line demarcates the lymph gland.

**Supplementary Figure 5. Ire1/Xbp1-dependent UPR pathway functions downstream of S-NO.**

**A-B.** Expression of *Xbp1-EGFP*, an ER stress marker, in the lymph gland (*Ubi-Gal4 UAS-Xbp1-EGFP*). The majority of *Xbp1-EGFP* is observed in the progenitor cell (left). A magnified image of the white dotted box (inset) is shown on the right. Low-level col^+^ (magenta) progenitors display *Xbp1-EGFP* (yellow arrow) **(A)**. A few Pxn^+^ (magenta) cells co-localize with *Xbp1-EGFP* (green) **(B)**. A magnified image of the white dotted box (inset) of the lymph gland is shown on the right.

**C-F**. Expression of *Drosophila* Atf4 (crc) protein in the lymph gland. Atf4 (magenta) and *Dome^Meso^*^+^ progenitors (green) are exclusive **(C)**. Compared to wild-type clones, clones expressing *Atf4* RNAi (*HHLT-Gal4 UAS-EGFP, UAS-Atf4 RNAi*; green) lack the expression of Atf4 (magenta). Magnified images are shown on the right. Feeding DTT, an ER stress inducer, enhances Atf4 protein levels (magenta) and nuclear localization (white arrow) in the lymph gland **(E)**. DAPI, blue. Atf6 protein (green) is observed in the salivary gland, which is increased by DTT feeding (left). DTT feeding does not modify the level of Atf6 in the lymph gland (right) (**F**).

**G-H.** Quantification of S-NO-dependent UPR gene expression in the lymph gland. Knockdown of *Nos* in *Dome^Meso^*^+^ progenitors reduces mRNA levels of *BiP* (Hsc70-3), *Xbp1*, *Pdi*, or *Ire1* (*Dome^Meso^-Gal4 UAS-EGFP, UAS-NosRNAi*)*.* Overexpression of *Nos* increases mRNA levels of *Xbp1, Pdi,* or *Ire1* mRNA (*Dome^Meso^-Gal4 UAS-EGFP, UAS-Nos)* **(G)***.* Overexpression of EGFR^C311S^ (*Dome^Meso^-Gal4 UAS-EGFP, UAS-FLAG::EGFR^C311S^*) in *Dome^Meso+^* progenitors increases the levels of *BiP* (Hsc70-3), *Xbp1*, *Pdi*, or *Ire1*, compared to controls or wild-type EGFR (*Dome^Meso^-Gal4 UAS-EGFP, UAS-FLAG::EGFR^WT^*) **(H)**. Statistical analysis: two-way ANOVA test.

White scale bar, 30 μm; yellow scale bar, 3 μm. White dotted lines demarcate the lymph gland. n.s.: not significant (*p*>0.01); ****p*<0.0001. Bars in graphs: Standard deviation.

**Supplementary Figure 6. S-NO is required for the maintenance of blood progenitors**

**A-B.** Expression of NimC1^+^ mature plasmatocytes in *Nos* mutants. Compared to wild-type lymph glands, hetero-allelic combinations of *Nos* mutants (*NosΔ^All^/NosΔ^15^* or *Nos^1^/NosΔ^15^*) reduce the differentiation of NimC1^+^ mature plasmatocytes (magenta) **(A)**. Quantification of NimC1^+^ mature plasmatocytes **(B)**. *Oregon R* (n=18), *NosΔ^All^/NosΔ^15^*(n=19)*, Nos^1^/NosΔ^15^* (n=14).

**C-D.** Reconstruction of differentiating progenitor cells expressing both *Dome^Meso^*^+^ and Pxn^+^ (yellow). Compared to wild-type lymph glands, lymph glands expressing *Nos^Oxy^* *RNAi* (*Dome^Meso^-Gal4 UAS-EGFP UAS-Nos^Oxy^ RNAi*) lead to a significant increase in the proportion of intermediate progenitors expressing both markers **(C)**. Quantification of differentiating progenitor cells (intermediate progenitors, IP) **(D)**. *Oregon R* (n=23), *Nos^Red^ RNAi* (n=24), *Nos^Oxy^ RNAi* (n=23).

**E-F.** Dome^Meso+^ progenitor-specific knockdown of *Nos* in the lymph gland. Compared to wild-type controls (left), *Nos^Oxy^* RNAi (BL28792) or *Nos^Red^* RNAi (VDRC27722) in *Dome^Meso^*^+^ progenitors does not alter the expression of NimC1^+^ mature plasmatocytes (magenta) **(E)**. Quantification of NimC1-expressing cells **(F)**. *Oregon R* (n=11), *Nos^Red^* RNAi (n=10), *Nos^Oxy^ RNAi* (n=16).

**G-H.** *Tep4*^+^ progenitor-specific knockdown of *Nos* in the lymph gland. Compared to wild-type controls (left), *Nos^Red^* RNAi or *Nos^Oxy^* RNAi does not change the expression of NimC1^+^ mature plasmatocytes (magenta) **(G)**. Quantification of NimC1^+^ plasmatocytes **(H)**. *Oregon R* (n=11), *Nos^Red^ RNAi* (n=13), *Nos^Oxy^ RNAi* (n=11).

**I-J.** Expression of RNAi against *Nos^Oxy^* in the intermediate progenitor does not alter the level of Pxn^+^ differentiating blood cells (*Nplp2-Gal4 UAS-EGFP UAS-Nos RNAi*) **(I)**. Quantification of Pxn^+^ area **(J)**. *Oregon R* (n=23), *Nos^Oxy^ RNAi* (n=20)*.*

**K-M.** Reducing *IP3R* expression in *Dome^Meso^*^+^ progenitors decreases *Dome^Meso+^* progenitor cells (green) and increases Pxn^+^ differentiating blood cells (magenta) **(K)**. Quantification of *Dome^Meso^*^+^ progenitors **(L)** and Pxn^+^ differentiating blood cells **(M)**. *Oregon R* (n=17), *IP3R RNAi* (n=8).

White scale bar, 30 μm; yellow scale bar, 3 μm. White dotted lines demarcate the lymph gland. n.s.: not significant (*p*>0.01); **p*<0.01; ****p*<0.0001. Bars in graphs: the median

**Supplementary Figure 7. Ire1/Xbp1-dependent UPR pathway is required for the progenitor maintenance**

**A-B.** UPR pathway genes do not modify the differentiation of mature plasmatocytes. Knockdown of the *Ire1*/*Xbp1* pathway (*Dome^Meso^-Gal4 UAS-EGFP UAS-Ire1 RNAi or UAS-Xbp1 RNAi*), the PERK/Atf4 pathway (*Dome^Meso^-Gal4 UAS-EGFP UAS-Pek RNAi or UAS-Atf4 RNAi*), or the Atf6 pathway (*Dome^Meso^-Gal4 UAS-EGFP UAS-Atf6 RNAi*) does not lead to changes in the proportion of NimC1^+^ plasmatocytes **(A)**. Quantification of NimC1^+^ cells per one lymph gland lobe **(B)**. *Oregon R* (n=18), *Ire1 RNAi* (n=14), *Xbp1 RNAi* (n=20), *Pek RNAi* (n=17*), Atf4 RNAi* (n=15), *Atf6 RNAi* (n=19).

**C-E.** *Ire1*/*Xbp1*-dependent UPR pathway maintains *Tep4*^+^ core progenitors. Expression of RNAi against *Ire1* or *Xbp1* (*Tep4-Gal4 UAS-EGFP UAS-Ire1 RNAi or UAS-Xbp1 RNAi*) decreases the proportion of *Tep4*^+^ core progenitors and increases Pxn^+^ differentiating blood cells **(C)**. Quantification of *Tep4*^+^ core progenitors **(D)** and Pxn^+^ differentiating blood cells per one lymph gland lobe **(E)**. *Oregon R* (n=17), *Ire1 RNAi* (n=18), *Xbp1 RNAi* (n=20).

**F-H.** PERK/Atf4 or Atf6 is not required for progenitor control. Knockdown of the *Pek*/*Atf4* pathway (*Dome^Meso^-Gal4 UAS-EGFP UAS-Pek RNAi or UAS-Atf4 RNAi*) or *Atf6* pathway (*Dome^Meso^-Gal4 UAS-EGFP UAS-Atf6 RNAi*) does not alter the proportion of *Dome^Meso+^* progenitors or Pxn^+^ differentiating blood cells **(F)**. Quantification of *Dome^Meso+^* area **(G)** and Pxn^+^ area **(H)**. *Oregon R* (n=18), *Pek RNAi* (n=17*), Atf4 RNAi* (n=15), *Atf6 RNAi* (n=19).

**I-K.** PERK/Atf4 or Atf6 is dispensable for the progenitor maintenance. Knockdown of the *Pek*/*Atf4* pathway (*Tep4-Gal4 UAS-EGFP UAS-Pek RNAi or UAS-crc RNAi*) or *Atf6* pathway (*Tep4-Gal4 UAS-EGFP UAS-Atf6 RNAi*) does not change *Tep4^+^* progenitors or Pxn^+^ differentiating blood cells **(I)**. Quantification of *Tep4* **(J)** and Pxn **(K)**. *Oregon R* (n=17), *Pek RNAi* (n=15*), Atf4 RNAi* (n=13), *Atf6 RNAi* (n=22).

White scale bar, 30 μm; yellow scale bar, 3 μm. White dotted lines demarcate the lymph gland n.s: not significant (p>0.01); ****p*<0.0001. Bars in graphs: the median

**Supplementary Figure 8. S-NO and UPR control G2 cell cycle arrest in blood progenitor cells**

**A-B.** S-NO and the Ire1/Xbp1 pathway control G2 cell cycle arrest in the lymph gland. Quantification of the number **(A)** or percentage **(B)** of *Dome^Meso+^* progenitors in each cell cycle (G2, yellow; G1, green; S, red) indicated by Fly-FUCCI in Figure 7.

**C-F.** Contribution of NO and cytosolic calcium to G2 cell cycle control. Reduction of *IP3R* expression inhibits G2 phase expression (white) in blood progenitor cells (*Dome^Meso^-Gal4 UAS-FUCCI UAS-IP3RRNAi*), which is not restored by simultaneous expression of *Nos* with *IP3R* RNAi (*Dome^Meso^-Gal4 UAS-FUCCI UAS-Nos UAS-IP3R RNAi*) or *Nos* RNAi with *IP3R* (*Dome^Meso^-Gal4 UAS-FUCCI UAS-IP3R RNAi UAS-Nos*). Increase of cytosolic calcium in blood progenitors does not alter the G2 cell cycle (*Dome^Meso^-Gal4 UAS-FUCCI UAS-IP3R*) **(C)**. Quantification of the number **(D)** of *Dome^Meso^*^+^ progenitors in the G2. *Oregon R* (n=36), *IP3R RNAi* (n=27), *UAS-Nos* (n=20), *UAS-IP3R* (n=24), *IP3R RNAi, UAS-Nos* (n=19)*, UAS-IP3R, NosRNAi* (n=19)*.* Quantification of the number **(A)** or percentage **(B)** of *Dome^Meso+^* progenitors in each cell cycle (G2, yellow; G1, green; S, red) indicated by Fly-FUCCI in Sup. Fig. 8C.

White scale bar, 30 μm; yellow scale bar, 3 μm. White dotted lines demarcate the lymph gland. n.s: not significant (p>0.01); ****p*<0.0001. Bars in graph **(D)**: the median.
